# Targeting macrophages with CAR-T cells delays solid tumor progression and enhances anti-tumor immunity

**DOI:** 10.1101/2021.12.17.473184

**Authors:** Alfonso R. Sánchez-Paulete, Jaime Mateus-Tique, Gurkan Mollaoglu, Sebastian R. Nielsen, Adam Marks, Ashwitha Lakshmi, Luisanna Pia, Alessia Baccarini, Miriam Merad, Brian D. Brown

## Abstract

Tumor-associated macrophages (TAMs) are one of the most abundant cell types in many solid tumors and typically exert protumor effects. This has led to an interest in macrophage-depleting agents for cancer therapy, but approaches developed to date have had limited success in clinical trials. Here, we report the development of a strategy for TAM depletion in mouse solid tumor models using chimeric antigen receptor (CAR) T cells targeting the macrophage marker F4/80 (F4.CAR-T). F4.CAR-T cells effectively killed macrophages in vitro and in vivo without toxicity. When injected into mice bearing orthotopic lung tumors, F4.CAR-T cells infiltrated tumor lesions and delayed tumor growth comparably to PD1 blockade, and significantly extended mouse survival. Anti-tumor effects were mediated by F4.CAR-T-produced IFN-γ, which promoted upregulation of MHC molecules on cancer cells and tumor-infiltrating myeloid cells. Notably, F4.CAR-T promoted expansion of endogenous CD8 T cells specific for tumor-associated antigens and led to immune editing of highly antigenic tumor cell clones. Antitumor impact was also observed in mouse models of ovarian and pancreatic cancer. These studies provide proof-of-principle evidence to support CAR-T targeting of TAMs as a means to enhance antitumor immunity.

## INTRODUCTION

Cancer immunotherapy is revolutionizing the treatment of many cancers (1). However, most patients with solid tumors do not respond to current immunotherapy agents. Among the factors involved in resistance to therapy are the tumor associate macrophages (TAM) (2). TAMs are one of the most abundant cell types in the tumor microenvironment (TME) of solid tumors (3) and adopt specific molecular states which are associated with poor clinical outcome (4). TAMs contribute to the suppression of antitumor immune responses and support tumor growth through a variety of mechanisms, including promoting angiogenesis, providing an immunosuppressive environment, helping to create a barrier that excludes effector T cells from the tumor core, capturing tumor antigens and preventing cross-presentation by cDC1, and promoting tumor invasiveness and metastasis (5).

The contribution of TAMs to the immunosuppressive TME has led to an interest in establishing clinically applicable strategies for depleting or reprogramming TAMs in human tumors (6). One prominent approach has been to target macrophage recruitment and survival by blocking the interaction between CSF1R on macrophages, and its ligand CSF1. The CSF1/R axis is critical to the differentiation and survival of macrophages, and inhibitors of this axis have shown reduction of macrophage density in solid tumors in mice and cancer patients (7). Despite this, CSF1/R inhibition has not been successful in the clinic as a monotherapy (8). More recently, strategies have been developed to modulate specific macrophage subtypes, such as those expressing TREM2, which are enriched among TAMs and correlate with T-cell exclusion (9,10). Efforts are also being pursued to reprogram TAMs from immunosuppressive to immunostimulatory, including through the use of TLR agonists and mRNA nanoparticles (6,11). These strategies are promising but have not been tested in the clinic yet, and thus there is still an important need to develop clinically applicable strategies to target TAMs and deplete or reprogram them.

One potentially promising, but still little explored, approach for TAM depletion is through the use of chimeric antigen receptor (CAR) T or NK cells (12–14). CAR-T cells are genetically modified to express an immunoglobulin-based receptor cognate to a specific surface antigen that triggers T-cell activation in a TCR-independent manner upon contact with an antigen-expressing cell (15). CAR-T cells have revolutionized the treatment of hematological malignancies, such as acute lymphoid leukemia (ALL) (16) and multiple myeloma (17). In most cases, the CAR-T target a marker specific to the lineage the malignant cells derive from. For example, CAR-T used in ALL target CD19, a molecule expressed by all normal and leukemic B cells. Continued innovation in the CAR-T field, including the implementation of CRISPR/Cas9 technology (18), makes it one of the most active areas of development in cancer immunotherapy (19).

Unfortunately, CAR-T therapy has been less successful for the treatment of solid tumors due to a number of issues (20). One of the most significant is the paucity of tumor-specific CAR-targetable antigens. There have been CAR designed against a variety of targets expressed by the cancer cells within solid tumors, such as ERB2, EGFR, and HER2, but none of these are specific to cancer cells, nor uniformly expressed across patients. Though innovative strategies are being developed to help restrict CAR-T cell killing to malignant cells (19), the cancer cells’ high capacity for adaption through gene mutations and epigenetic silencing often leads to loss or reduction of target expression on the cancer cells, enabling them to evade CAR-T killing. Additional to the problem of antigen availability is the challenge for the CAR-T to overcome the highly immunosuppressive and immune excluded environment of solid tumors, which, as noted, is facilitated by TAMs.

Here we set out to develop and evaluate CAR-T cells targeting macrophages as a means to achieve antitumor impact and reprogram the immunosuppressive TME of solid organ tumors. The rational for using CAR-T to target macrophages is to eliminate the immunosuppressive activity of TAMs, while simultaneously achieving local release of pro-inflammatory cytokines, such as IFN-γ and TNF, within tumors, which may facilitate endogenous innate and adaptive immune responses. To this end, we generated a 2^nd^ generation CAR targeting the mouse pan-macrophage marker F4/80 (F4.CAR) and show that T cells expressing F4.CAR (F4.CAR-T) specifically killed F4/80+ macrophages and eosinophils. Injection of F4.CAR-T into immunocompetent mice bearing lung tumors resulted in slower tumor growth and prolonged survival, despite cancer cells not being directly targeted by treatment. Antitumor benefit was also observed in models of ovarian and pancreatic cancer. Notably, we established that treatment promoted expansion of endogenous CD8+ T cells specific for tumor antigen and immune editing of the tumor, and this was dependent on the F4.CAR-T expressing IFN-γ. These studies help to establish macrophage-targeted CAR-T cells as a means to potentiate endogenous antitumor immunity and slow tumor growth.

## RESULTS

### F4.CAR-T cells recognize and kill mouse macrophages

We set out to generate a CAR with specific tropism for macrophages. We aimed to target macrophages broadly as a proof-of-principle for this approach. We chose to target the Adgre1 gene product, F4/80, which is highly and specifically expressed by mouse macrophages, as well as eosinophils and certain populations of monocytes (21). We sequenced the immunoglobulin variable domains of the F4/80 hybridoma (22) and cloned the heavy and light chain fractions of the variable region as a single-chain fragment variable (ScFv) into a CD28/CD3z-based CAR retroviral vector (23). A GFP moiety, separated from the CAR by a T2A self-cleaving peptide, was cloned downstream of the CAR coding region to mark transduced cells (**Fig 1A**). We isolated CD8^+^ T cells from wildtype mice and transduced them with the F4.CAR vector or a control retroviral vector encoding GFP. Expression of the CAR and GFP by transduced T cells was confirmed by flow cytometry (**Fig 1B**).

**Figure 1.**
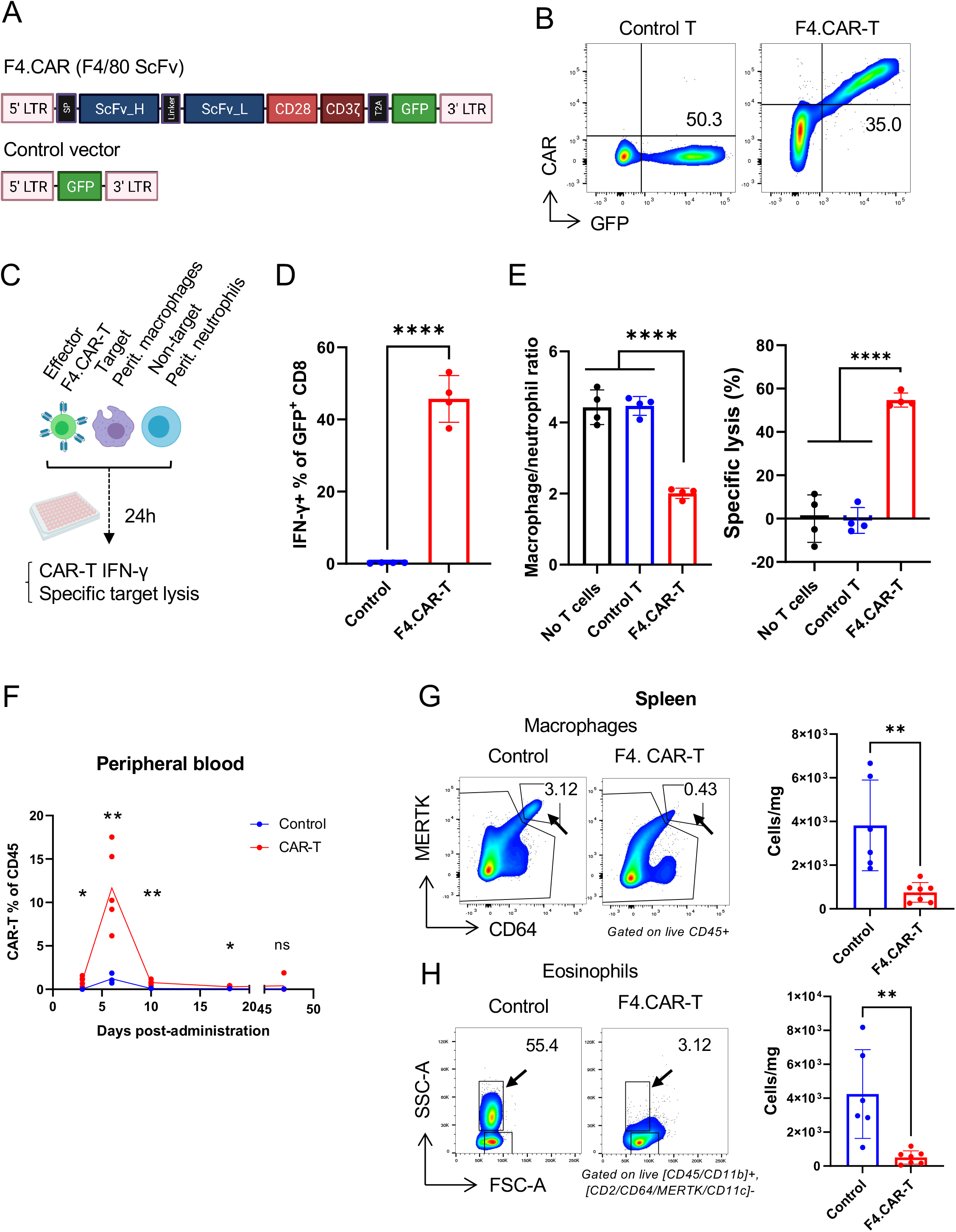
F4.CAR-T cells recognize and specifically lyse mouse macrophages. **(A)** Schematic depicting the structure of the F4.CAR construct and control vector for T-cell transduction. **(B)** Representative flow cytometry plots showing expression of the CAR and GFP in control or transduced CD8 T cells. Shown are the percentages of transduced cells. **(C-E)** Ex vivo CAR-T killing assay. Mice were i.p. injected with 1.5 ml 3% thioglycolate broth and peritoneal lavage obtained 72h later and cocultured with control or F4.CAR-T cells for 24h. **(C)** Schematic explaining the assay design. **(D)** Percentage of GFP+ T cells stained for intracellular IFN-γ at the end of the coculture. **(E)** The ratio of peritoneal macrophages to neutrophils in the presence of control or F4.CAR-T cells is shown on the left. On the right, specific lysis of macrophages was calculated using neutrophils as non-targeted cells. *p<0.05, **p<0.01, ***p<0.001, ****p p<0.0001 **(F)** Percentage of CAR-T cells among peripheral blood cells over time after i.v. CAR-T cell administration. **(G,H)** On the left, representative flow cytometry plots from spleens of control- and CAR-T mice treated showing the macrophage **(G)** and eosinophil **(H)** populations. Numbers represent percentage of parent populations. On the right, quantification of macrophages (G) and eosinophils (H) per milligram of spleen. Complete gating strategies can be found in supplementary figure 2. SP: signal peptide; LTR: long terminal repeat; E:T: effector:target; i.v.: intravenous. *p<0.05, **p<0.01, ***p<0.001 (t-student)

To test the activity of F4/80 CAR-T cells (F4.CAR-T), we cocultured them with primary macrophages ex vivo. To this end, we generated peritoneal thioglycolate-elicited macrophages and co-cultured them with F4.CAR-T CD8 T cells to interrogate target-specific lysis. Peritoneal neutrophils acted as non-target internal controls (**Fig 1C**). GFP+ F4.CAR-T but not control T cells produced IFN-γ at abundant levels in this setting (**Fig 1D**). This was accompanied by the specific lysis of macrophages (**Fig 1E**). We confirmed the specificity of F4.CAR-T activity using RAW264.7 cells, a mouse macrophage cell line which constitutively expresses F4/80. In this setting we used A20 cells, an F4/80-negative B cell line, as an internal control for the quantification of specific lysis (**Supp Fig 1A**). Similar to peritoneal macrophages, RAW264.7 induced IFN-γ production by F4.CAR-T cells (**Supp Fig 1B**) and were specifically killed by these cells (**Suppl Fig 1C**). These results indicate that the F4.CAR construct is functional and facilitates lysis of macrophages by F4.CAR-expressing CD8 T cells.

We next evaluated F4.CAR-T activity in vivo. While CAR-T injection is normally preceded by lymphodepleting conditioning to facilitate CAR-T expansion, we reasoned that F4.CAR-T may not require conditioning because of the ample availability of target cells (macrophages) and their nature as antigen-presenting cells. To test this, we injected 5×10^6^ F4.CAR-T i.v. into syngeneic healthy C57Bl/6 mice and measured expansion in peripheral blood by flow cytometry detection of GFP+ cells. As a control, we injected mice with T cells transduced with a retroviral vector encoding GFP. By 6 days, F4.CAR-T reached 5-15% of circulating CD45+ cells and contracted shortly afterwards, whereas control T cells did not expand (**Fig 1F**). To evaluate F4.CAR-T functionality, we collected the spleens from the mice, stained for a panel of immune cell markers and analyzed by flow cytometry. In mice injected with control T cells, the frequency of macrophages in the spleen averaged 4×10^3^ cells per mg of tissue (**Fig 1G**). In contrast, in mice that received the F4.CAR-T, macrophage frequency was reduced by 80%. Eosinophils, which express F4/80, were similarly eliminated in the F4.CAR-T treated mice (**Fig 1H**). Mice receiving F4.CAR-T cells did not show signs of stress and no lethality was observed from treatment. These results indicate that F4.CAR-T cells were effective in target cell killing in vivo.

### F4.CAR-T cells kill TAMs in HKP1 lung tumors

Macrophages are abundant in non-small cell lung carcinoma (NSCLC) lesions and considerable evidence indicates that they play a key role in promoting immunosuppression in the TME. To model NSCLC in mice, we utilized HKP1 cells (24), which carry an activating KRAS allele (Kras^G12D^) and a deletion of p53 (p53^-/-^), both common genetic mutations in human NSCLC. HKP1 also express luciferase and tdTomato, which enables the tumors to be tracked in vivo. HKP1 were injected i.v. to seed lung tumors. At 19 days post-injection, there was abundant macrophage accumulation in the tumor lesions (**Fig 2A**). We performed flow cytometry analysis of healthy and tumor-bearing lungs and observed that, in tumor-bearing lungs, monocyte-derived macrophages (Mo-Macs) were recruited to lungs and outnumbered resident alveolar macrophages (AM) (**Fig 2B, Supp. Fig 2**), fitting with previous findings in human cancer lesions and mouse models (25,26). We quantified tumor burden histologically by calculating the percentage of tumor tissue area in lung H&E-stained sections. The total number of Mo-Macs in the lungs of tumor-bearing mice was proportional to histologically calculated tumor burden, underlining the association between tumor growth and macrophage accumulation (**Fig 2C**).

**Figure 2.**
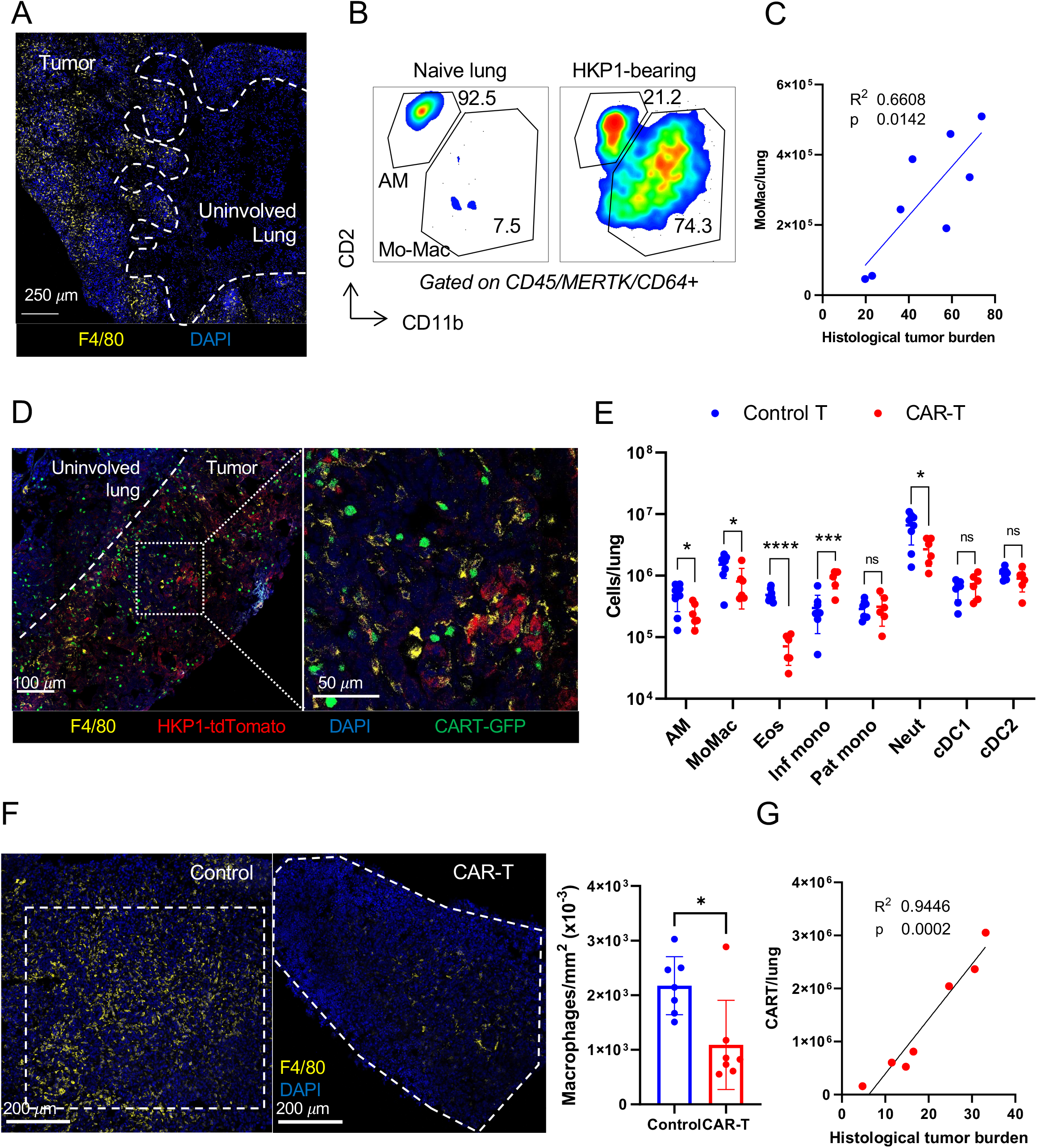
F4.CAR-T cells successfully infiltrate HKP1 NSCLC tumors and eliminate tumor-associated macrophages. HKP1 tumors were established and CAR-T cells administered 7 (A, B, D, F) or 12 (C, E, G) days later. Lung samples were obtained 19 days after tumor inoculation and processed for flow cytometry or immunofluorescent tissue staining. **(A)** Representative confocal microscopy sample of macrophage infiltration in an HKP1 lung tumor. Dashed line demarcates the tumor lesion. Color coding is indicated at the bottom. **(B)** Representative flow cytometry plots of lung macrophages from healthy and HKP1-bearing lungs. Numbers denote percentage of macrophages contributed by each subpopulation. **(C)** Correlation between the number of monocyte-derived macrophages per lung and histological HKP1 tumor burden (calculated as described in Materials and Methods). **(D)** Confocal microscopy section of HKP1 lesions in a mouse lung. Right panel represents a zoom in of the area dotted in the left image. Dashed line demarcates the tumor lesion. Color coding is indicated at the bottom. **(E)** Immune cell populations in lungs of control and CART-treated mice. Complete gating strategies can be found in supplementary figure 2. Representative experiment of five is shown. **(F)** Representative regions within HKP1 tumors were defined and the number of macrophages in those regions calculated by F4/80 immunofluorescent staining using QuPath software. Left panel shows representative samples from control and CART-treated tumors, with dashed lines defining the studied regions. Right panel shows the quantification of macrophages per square mm within these regions. **(G)** Correlation between the number of CAR-T cells per lung and histological HKP1 tumor burden. AM: alveolar macrophage; Mo-Mac: monocyte-derived macrophage; Eos: eosinophil; Inf mono: inflammatory monocyte; Pat mono: patrolling monocyte; Neut: neutrophil; cDC1/2: conventional dendritic cell type 1/2. *p<0.05, **p<0.01, ***p<0.001, ****p p<0.0001 (t-student)

We next set out to determine if F4.CAR-T could deplete the macrophages in the HKP1 tumor lesions. Seven days after HKP1 injection, we treated mice with control T cells or F4.CAR-T cells, and 12 days later we collected lungs and examined tissue by immunofluorescence confocal microscopy and flow cytometry. Microscopy analysis showed F4.CAR-T highly and preferentially infiltrated lung tumor lesions and co-localized with F4/80+ cells at tumor sites (**Fig 2D**). Importantly, there was a substantial reduction in the number of macrophages and eosinophils in the tumor-bearing lungs of F4.CAR-T treated animals compared to controls (**Fig 2E**), whereas the number of DCs and patrolling monocytes remained the same. We also observed an increase in the number of inflammatory monocytes in the lungs, which could reflect an ongoing homeostatic response to replenish the depleted macrophage population.

A limitation of flow cytometry in the study of orthotopic lung tumors is the inability to parse cells originating from healthy and tumorous tissue. To verify if depletion of target cells was occurring in the tumor lesions, we generated lung tissue sections from control and F4.CAR-T treated mice bearing HKP1 tumors and analyzed defined tumor regions to quantify the number of F4/80+ cells per unit of area within tumors. This analysis showed that the density of macrophages in the TME of F4.CAR-T-treated mice was markedly reduced across specimens, to approximately half of the control-treated counterparts (**Fig 2F**).

Flow cytometry of single-cell suspensions from the same lungs indicated that the total number of CAR-T cells detected per lung was directly proportional to the histological tumor burden in those mice (**Fig 2G**). This is most likely related to the accumulation of macrophages during tumor growth (**Fig 2C**), and it corroborates the observed preference for F4.CAR-T cells to localize to tumor sites.

Together, these results demonstrate that F4.CAR-T were able to infiltrate lung tumor lesions, eliminate macrophages in the TME, and reduce TAM density in lung tumors.

### F4.CAR-T cells delay growth of HKP1 lung tumors and extend survival of tumor-bearing mice

We next set out to assess whether F4.CAR-T administration could impact HKP1 tumor growth. We injected mice i.v. with HKP1 cells, followed 7 days later by injection of control T cells or F4.CAR-T cells (**Fig. 3A**). We monitored tumor growth non-invasively by luciferase-based luminescence using an IVIS system. We could detect comparable tumors prior to and 3 days post-treatment, but within 10 days of treatment, we found a 51% reduction in tumor burden in mice that received F4.CAR-T, as measured by photon detection (**Fig. 3B)**.

**Figure 3.**
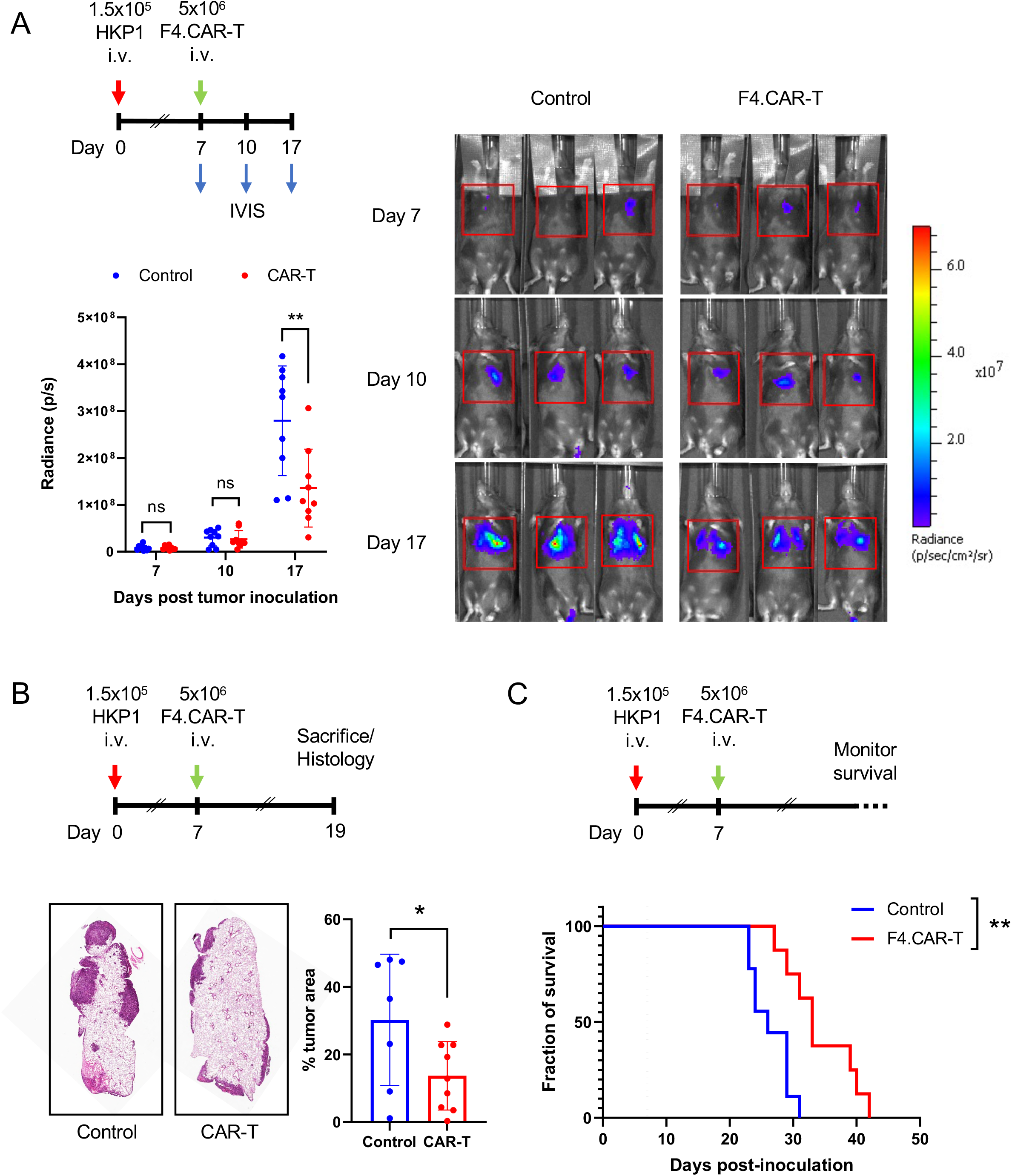
F4.CAR-T cells delay growth of HKP1 lung tumors and extend the survival of tumor-bearing mice. HKP1 tumors were established and CAR-T cells administered according to the treatment schematic on each panel. **(A)** Bioluminiscence readings from the thoracic area (red square on the images) were taken at the time points indicated. Right panel shows representative IVIS images from control and CART-treated mice and color scale for radiance. Left bottom panel shows quantification of luciferase radiance signal from control and CART-treated mice. A representative experiment of three performed is shown. **(B)** Mice were euthanized at the indicated time point and lungs processed for H&E staining. Left panel shows representative sections from control or F4.CAR-T treated mice. Histological tumor burden was calculated as described in materials and methods. A representative experiment of >5 performed is shown. **(C)** HKP1-bearing mice were followed for survival after CAR-T treatment. A representative experiment of three performed is shown. *p<0.05, **p<0.01, ***p<0.001

In successive experiments, we sacrificed mice 19 days after tumor cell inoculation, extracted the lungs and performed H&E staining to quantify tumor burden by histological analysis (**Fig. 3C)**. We consistently observed significantly reduced tumor burden in F4.CAR-T-treated mice compared to mice injected with control T cells (**Fig 3D**). Importantly, this delay in tumor growth translated into a survival benefit for treated mice, increasing the median survival by 28% from 23.5 to 30 days (**Fig 3E and 3F**). Similar results were obtained when mild lymphodepletion (2 Gy total-body irradiation) was used prior to CAR-T administration (**Suppl Fig 3A)** These results show that in vivo targeting of macrophages by F4.CAR-T cells has an antitumor effect in the HKP1 model of NSCLC.

### CAR-T targeting of TAMs induces tumor antigen-specific T-cells and immune editing of tumors

Though tumor burden was significantly reduced, we were still able to detect tumors in F4.CAR-T-treated mice. When we examined these tumors by florescence microscopy, we unexpectedly observed that there was a major reduction in tdTomato detection in HKP1 tumors from F4.CAR-T treated mice (**Fig. 4A**). Using flow cytometry, we quantified the mean florescence intensity (MFI) for tdTomato in HKP1 cells and confirmed a strong reduction in F4.CAR-T-treated mice (**Fig. 4B**). We also found a direct correlation between the level of expression of tdTomato and tumor burden in untreated mice: mice with higher tumor burden displayed a higher per-cell expression of tdTomato (**Fig 4C**), suggesting that naturally occurring pressure against tdTomato in this tumor model can delay its development.

**Figure 4.**
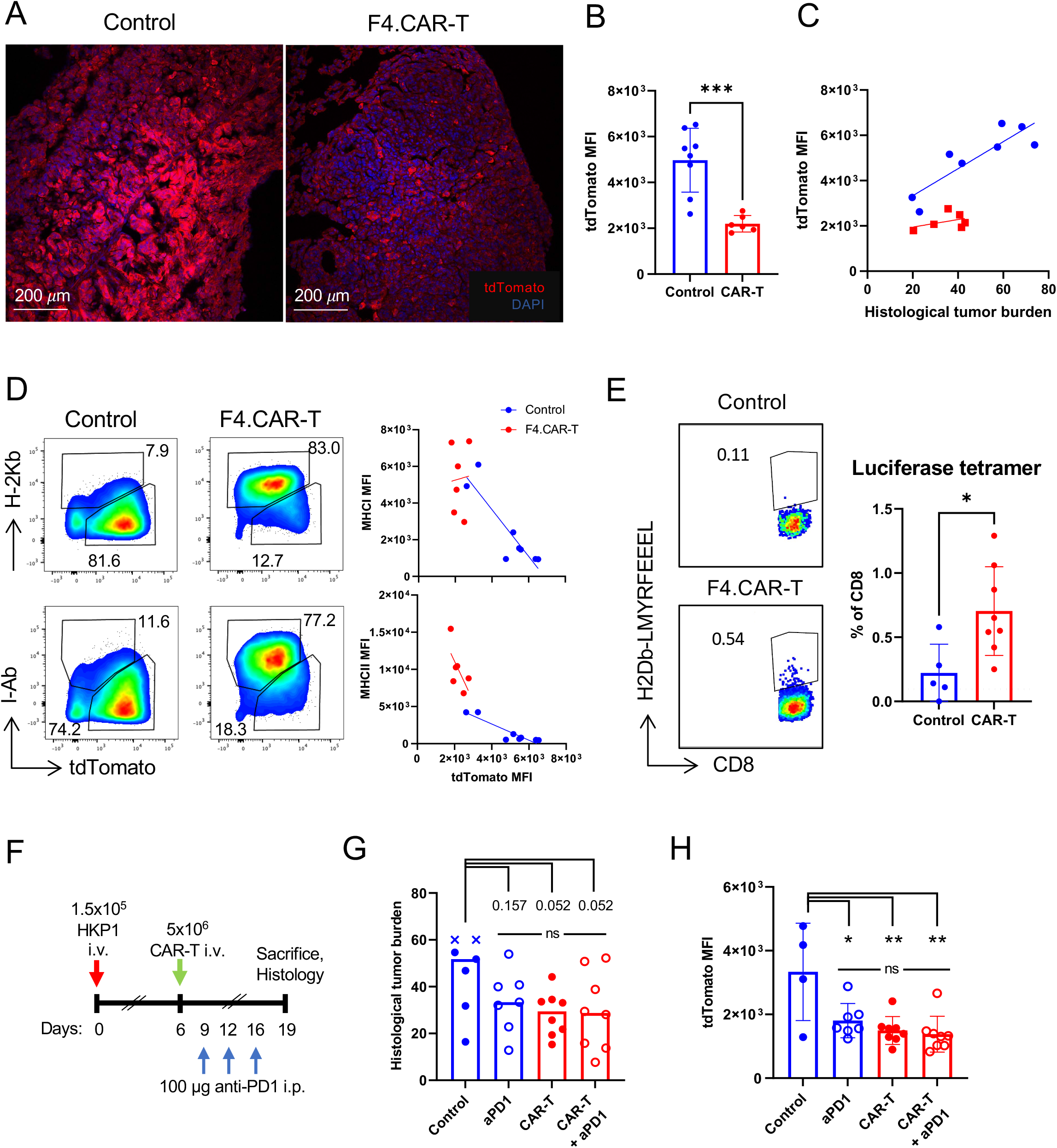
F4.CAR-T cells promote tumor immune editing and expansion of tumor-antigen specific T cells. (A-D) HKP1 tumors were established and CAR-T cells administered 12 days afterwards. At day 19, lung samples were processed for confocal microscopy or flow cytometry. **(A)** Shown is tdTomato detection in representative HKP1 tumors from control or CART-treated mice by confocal microscopy. **(B)** Flow cytometry was used to quantify tdTomato MFI in HKP1 cells from control and CART-treated mice. Results are representative from >5 experiments performed. **(C)** Correlation of HKP1 tdTomato MFI with histological tumor burden. **(D)** Left shows representative dot plots of HKP1 cells from control and CART-treated mice showing tdTomato vs MHC-I or MHC-II detection. Right graphs show the correlation between tdTomato and MHC-I or MHC-II MFI from HKP1 cells in control and CART-treated mice. **(E)** HKP1 tumors were established and CAR-T cells administered 7 days afterwards. At day 19, lung single-cell suspensions were stained with H2Db-LMYRFEEEL tetramers as detailed in Materials and Methods. Left panels show representative dot plots for tetramer staining and percentage of positive cells within the endogenous CD8 T cell population. Right panel shows percentages of tetramer^+^ CD8 T cells across samples. One experiment representative of three performed. **(F)** HKP1 tumors were established and treated with F4.CAR-T cells and/or anti-PD1. **(G)** Histological tumor burden in control and treated mice. One experiment representative of three performed. **(H)** tdTomato MFI in HKP1 tumor cells in control and treated mice, calculated by flow cytometry. *p<0.05, **p<0.01, ***p<0.001, ****p p<0.0001

In previous studies, using another visualizable protein, GFP, as a cancer-associated antigen, we had observed an inverse correlation between MHC and GFP expression on cancer cells, which was indicative of tumor evolution against an antigen-directed T cell response (27). To assess whether similar dynamics may be occurring upon treatment with F4.CAR-T, we stained single cell suspensions of mouse lungs for MHC-I and MHC-II and analyzed by flow cytometry. We observed that HKP1 cells from high-tumor-burden control mice presented high tdTomato expression and little to no detectable MHC-I or MHC-II. In contrast, in F4.CAR-T treated mice and a fraction of low-tumor burden untreated mice, HKP1 cells showed starkly elevated levels of MHC-I and MHC-II, coinciding with reduced per-cell expression of tdTomato and reduced tumor burden (**Fig. 4D**). These observations were consistent across >5 independent experiments.

The reduction of tdTomato levels and increased MHC expression in tumor cells, associated with a reduced tumor burden phenotype, suggested F4.CAR-T treatment may be promoting an endogenous T cell response against tdTomato as a tumor-associated antigen. Though a tetramer is not available for tdTomato-specific T cells, one is available for an immunodominant epitope of luciferase [H2Db-LMYRFEEEL (28)], which is also expressed by HKP1 cells. We injected mice with HKP1 cells, followed by control or F4.CAR-T, as above, and after 19 days, collected lungs, made single cell suspensions, and stained with this tetramer to detect luciferase-specific T cells by flow cytometry. Consistent with our hypothesis, there was a significant increase in the frequency of luciferase-specific CD8 T cells in the F4.CAR-T treated mice compared to control animals (**Fig 4E**).

Taken together, these findings indicate treatment with F4.CAR-T not only eliminated tumor macrophages, but also promoted expansion of tumor-antigen associated T cells and immune editing of tumor cells. This helps provide insight into the mechanisms by which F4.CAR-T mediate reduction of tumor burden.

### F4.CAR-T reduces HKP1 lung tumor burden to a similar magnitude as PD1 blockade

HKP1 tumors have been previously shown to be partially responsive to PD1 blockade (29). We sought to assess how F4.CAR-T treatment compares to anti-PD1, and to evaluate whether they might synergize. We injected HKP1-bearing mice with F4.CAR-T, anti-PD1, or both anti-PD1 and F4.CAR-T (**Fig 4F**). As expected, anti-PD1 trended to reduce tumor burden. F4.CAR-T mediated a similar reduction, indicating equivalent efficacy of the two approaches. Combination treatment of anti-PD1 and F4.CAR-T led to a decrease in tumor burden, but it did to the same extent as each of the monotherapies (**Fig 4G**). When we analyzed lungs by flow cytometry, we once again found that treatment with F4.CAR-T led to a significant decrease in tdTomato^high^ HKP1 cells: however, we also discovered this same process was independently facilitated by PD1 blockade (**Fig 4H**), hinting at a shared mechanism of action between both strategies. Combined treatment did not further cause reduction in tdTomato expression by HKP1 cells.

These observations suggests that CART-mediated macrophage depletion and PD1 blockade both delayed HKP1 tumor growth by facilitating endogenous T cell-mediated elimination of the most highly antigenic tumor cells. This mechanistic redundancy could provide an explanation for the lack of synergy between both strategies in this tumor model.

### Expression of IFN-γ by F4.CAR-T cells mediates TME reprogramming and antitumor effects

Our observations that treatment with F4.CAR-T led to increased MHC class I and II expression on cells in the tumor led us to consider how this might be mediated. Amongst the strongest inducers of MHC is IFN-γ, which is one of the dominant cytokines produced by activated T cells. We thus hypothesized that F4.CAR-T-mediated release of IFN-γ may be playing a role in sensitizing the tumors to cellular immunity.

To determine if F4.CAR-T-produced IFN-γ was playing a significant role in tumor growth delay, we isolated CD8+ T cells from Ifng-/- mice, transduced them with the F4.CAR vector, and injected them into mice bearing HKP1 lung tumors (**Fig 5A**). As a comparison control, mice were injected with wildtype F4.CAR T cells. In mice injected with Ifng-/- F4.CAR-T cells, the delay in tumor growth mediated by wildtype F4.CAR-T was completely abrogated (**Fig 5B**). Furthermore, HKP1 tumor cells from mice treated with IFN-γ-deficient CAR-T failed to upregulate expression of MHC-I, MHC-II or PDL1 (**Fig 5C**). Coinciding with the absence of upregulated MHC-I and MHC-II, use of Ifng-/- CAR-T cells did not lead to a decrease in tdTomato^high^ cells, as occurred in F4.CAR-T treated mice (**Fig 5D**). This was not because IFN-γ directly dampens tdTomato expression, as in vitro treatment of HKP1 cells with IFN-γ did not affect tdTomato expression (**Suppl. Fig 4**).

**Figure 5.**
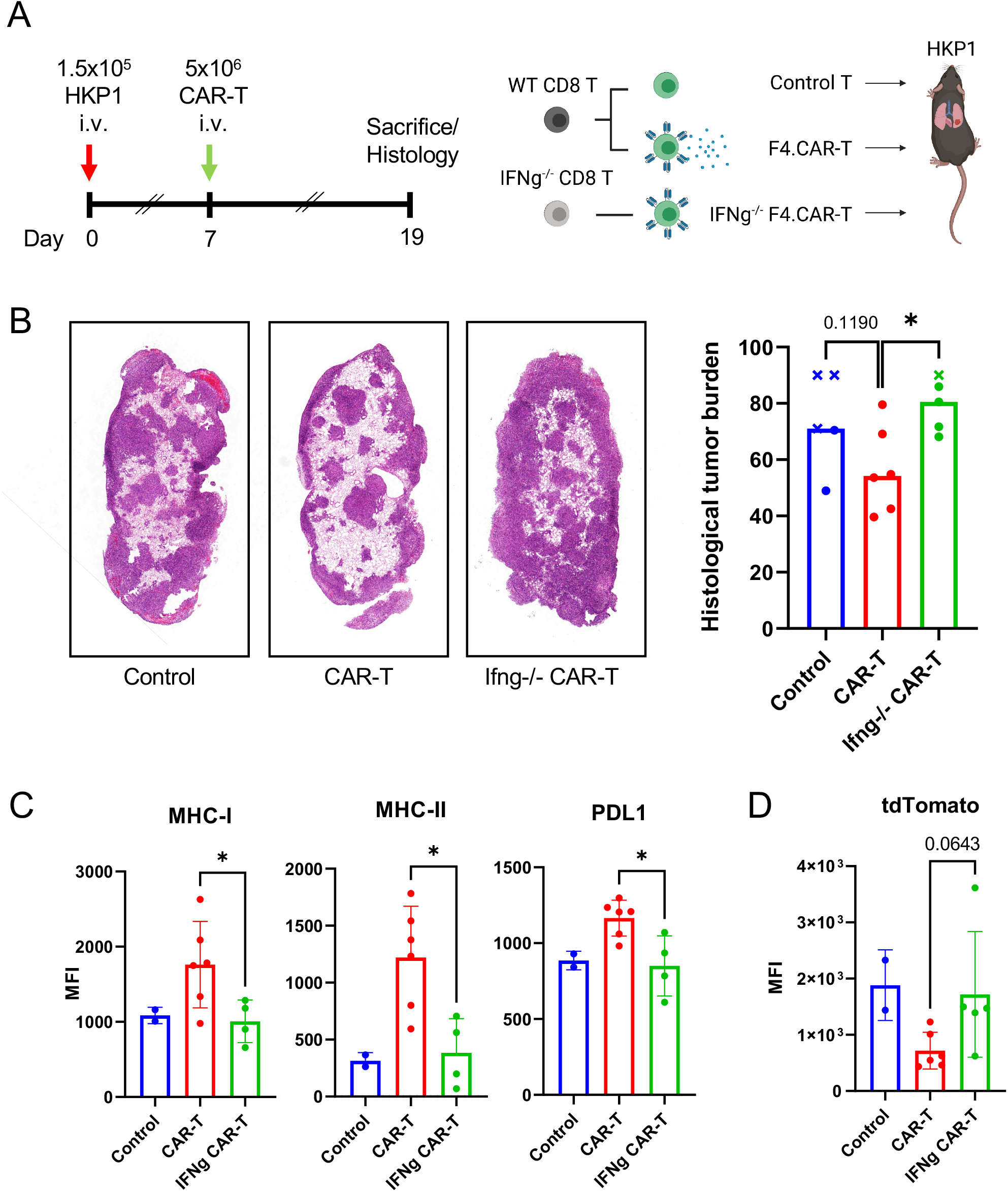
IFN-*γ* production by F4.CAR-T cells reprograms the HKP1 tumor microenvironment and is necessary for its antitumor effect. **(A)** Left shows experiment schematic. Right cartoon depicts F4.CAR-T generation from IFNg^-/-^ or WT mice. **(B)** Lungs were obtained and processed for histological analysis. Shown left are representative samples of H&E stainings from the indicated groups. On the right is shown the quantification of tumor burden across all mice, with X datapoint symbols indicating mice that died before the designed time of sacrifice. Columns represent median values. Mann-Whitney ranked analysis was used for statistics. **(C,D)** Shown are the mean fluorescent intensities for the indicated elements on HKP1 tumor cells in control and treated mice. *p<0.05, **p<0.01, ***p<0.001, ****p p<0.0001

Other immune cells in the TME were also found to show increased levels of MHC-II and PDL1 in F4.CAR-T treated mice (**Suppl. Fig 5A**). These changes were again dependent on IFN-γ from F4.CAR-T cells since upregulated MHC-II and PD-L1 was not observed in tumor-bearing mice injected with Ifng-/- F4.CAR-T (**Suppl. Fig 5B**). Ifng-/- F4.CAR-T were not defective in their ability to infiltrate tumor-bearing lungs, as they reached comparable numbers to their wildtype counterparts (**Suppl. Fig 5C**). Even more notably, Ifng-/- F4.CAR-T cells retained their ability to lyse target populations (monocyte-derived macrophages and eosinophils) to a level that was comparable to wildtype F4.CAR-T cells (**Supp Fig 5D**).

These studies indicate that F4.CAR-T-mediated control of tumor burden and immune editing is largely facilitated by release of IFN-γ, and that macrophage killing may not be necessary for treatment efficacy. Tumor macrophages may thus serve as a fixed target for CAR-T homing into the TME and release of IFN-γ.

### F4.CAR-T cells delay growth of ID8 ovarian tumors and KPC pancreatic tumors

A recent study employing a CAR targeting Folate Receptor beta (FRB)-expressing macrophages demonstrated delay of ID8 ovarian tumor growth in mice (14). To determine if F4.CAR-T could also be effective in this tumor setting, we evaluated their efficacy in an aggressive ID8 ovarian tumor model in which the ID8 overexpress VEGF-A and Defensin Beta 29 (30). When injected into the peritoneal cavity, ID8 cells form disseminated tumors in the omentum and on the diaphragm, which are abundantly surrounded by macrophages (31). In this model, tumor growth is associated with the development of peritoneal ascites. We injected mice with ID8 cells i.p. and treated them 2 weeks later with control T cells or F4.CAR-T (**Fig 6A**). Though tumors grew in both groups, treatment with F4.CAR-T cells delayed weight gain and the development of tumor-induced ascites in treated mice (**Fig 6B,C**).

**Figure 6.**
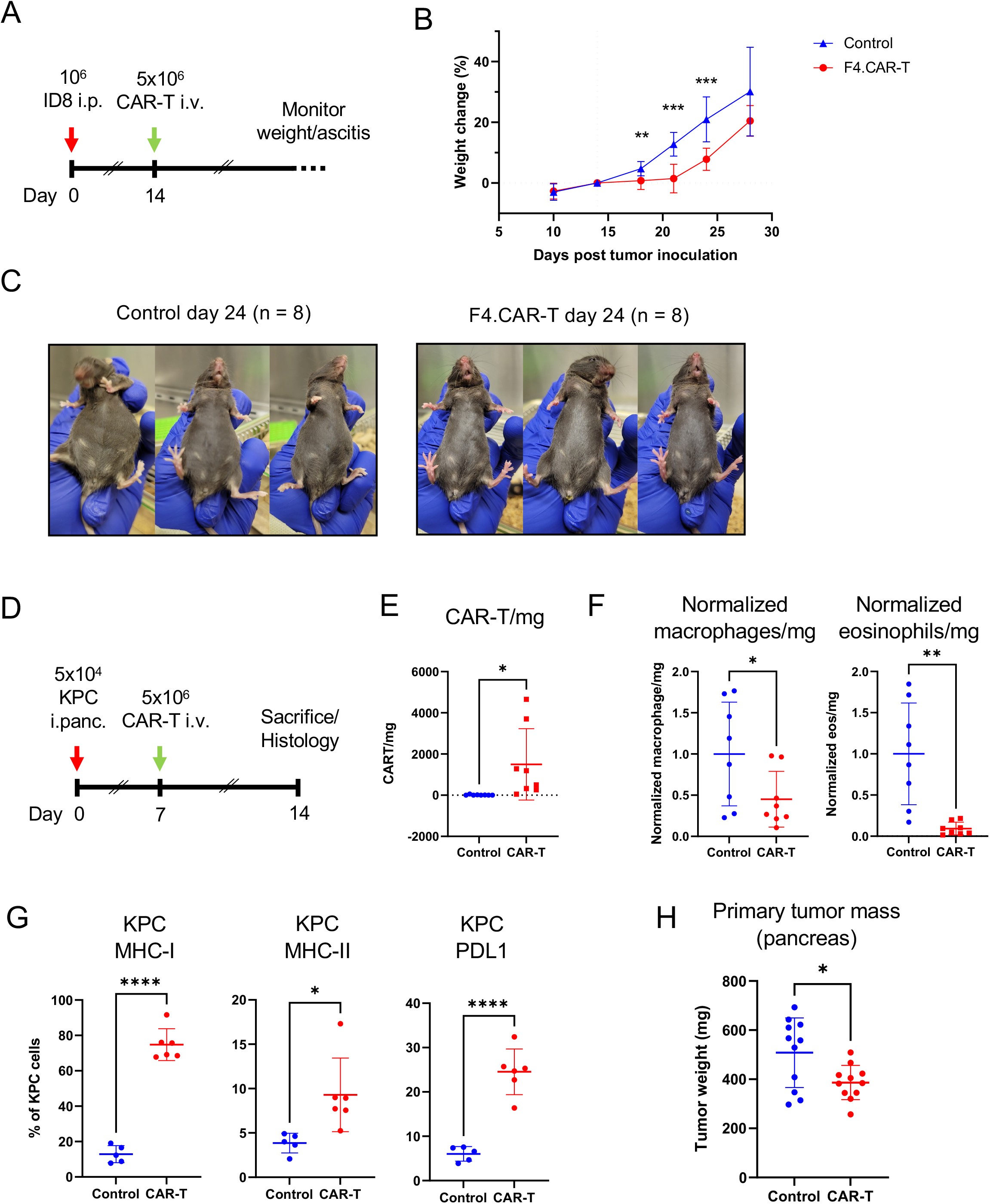
F4.CAR-T cells delay growth of ID8 ovarian and KPC pancreatic tumors. **(A-C)** ID8_VEGF tumors were established by i.p. injection of 10^6^ cells. 5×10^6^ F4.CAR-T cells were injected i.v. on day 14. Shown is one experiment, representative of two performed. **(A)** Experiment schematic. Mouse weight was obtained and mice monitored for development of ascites biweekly. **(B)** Change in body weight in control and CAR-T-treated mice, normalized to the time of treatment (day 14). **(C)** Representative examples of tumor-induced ascites at day 24 post-tumor inoculation. **(D-G)** KPC tumors were established by surgical injection of 5×10^4^ cells into the pancreas. 5×10^6^ F4.CAR-T cells were injected i.v. 7 days later. Mice were sacrificed on day 14 and pancreas tumors processed for flow cytometry. Results from two pooled experiments are shown for (E,F,H) **(D)** Experiment schematic. Shown are two pooled experiments (E,F,H). **(E)** Number of F4.CAR-T cells per mg of tissue in primary pancreas tumors. **(F)** The number of cells per mg of tumor from two pooled experiments was normalized to the control average for each experiment. Shown are normalized macrophages and eosinophils in control and F4.CAR-T treated mice. **(G)** MFI for MHC-I and PDL1 on KPC tumor cells. KPC cells were defined as DAPI/CD45-, FSC/SSChi, analogously to the strategy defined for HKP1 in Supp. Fig. 1. **(H)** Mass of primary pancreas tumors at day 14. *p<0.05, **p<0.01, ***p<0.001, ****p p<0.0001

We next evaluated F4.CAR-T cells activity in the context of pancreatic cancer, a typically macrophage-rich, T-cell excluded malignancy with few therapeutic options (32,33). We utilized the KPC pancreatic cancer model in which the cancer cells carry a Kras^G12D^ mutation and p53 deletion (34), which are two of the most common mutations in human pancreatic cancer (35). KPC cells were surgically inoculated into the pancreas of mice and control T cells or F4.CAR-T cells were injected i.v. 7 days afterwards. At day 14 after tumor inoculation, mice were euthanized and pancreas lesions weighed as a readout for antitumor effect and processed for flow cytometry and histology (**Fig 6D**). By flow cytometry, we observed F4.CAR-T cells successfully infiltrated tumor masses (**Fig 6E**), reduced the density of macrophages and eosinophils in the tumor lesions (**Fig 6F**) and led to upregulation of MHC-I and II on tumor cells, as well as PD-L1 (**Fig 6G**). Importantly, F4.CAR-T treatment reduced the mass of the primary pancreatic lesions at the time of sacrifice (**Fig 6H**). These studies demonstrate that F4.CAR-T can mediate anti-tumor effects in different tumor types.

## DISCUSSION

We show here that targeting macrophages with CAR T cells can delay tumor growth in mouse models of lung, ovarian and pancreatic cancer. Antitumor activity against NSCLC was associated with the expansion of tumor antigen-specific T cells and tumor immune editing, and was dependent on IFN-γ production by the CAR-T cells, which stimulated upregulation of antigen presentation by tumor and myeloid cells in the TME. Some of these mechanistic traits could be also observed in the ovarian and pancreatic tumor models.

Previous reports have explored CAR-T and -NK targeting of TAMs or MDSCs in mouse tumor models. Ruella et al. identified CD123 as a shared antigen between Hodgkin lymphoma cells and M2-like TAMs and showed that tumor-directed CD123 CAR-T cells could also be active against TAMs, avoiding immune suppression (12). Parihar et al. developed CAR-NK cells targeting NKG2D ligands expressed on tumor-associated myeloid-derived suppressor cells, achieving significant antitumor efficacy in a xenograft neuroblastoma model in mice (13). Most recently, Rodriguez-Garcia et al. described the use of macrophage-targeted CAR-T cells to treat ID8 ovarian tumors in mice (14). In this work, they used a CAR directed against Folate Receptor beta (FRB), which is upregulated on a subset of inflammatory macrophages, including many TAMs. They showed that FRB-targeted CAR-T cells depleted macrophages in the peritoneal cavity and extended the survival of ID8 tumor-bearing mice. Promisingly, both Parihar and Rodriguez-Garcia showed that combining myeloid-targeted CAR-T or -NK with tumor-targeting CAR-Ts synergistically enhanced mouse survival in the models tested (13,14). Here, we expand on these findings by showing that CAR-T targeting of macrophages can additionally delay tumor growth in models of NSCLC (HKP1 tumor model) and pancreatic cancer (KPC tumor model), and by providing mechanistic insight into the processes leading to tumor growth delay in these models.

When analyzing HKP1 tumor lesions, we observed a striking decrease in tdTomato levels within tumors that had persisted in F4.CAR-T treated animals. KP tumors, an established NSCLC model (36), are not very immunogenic, but the incorporation of tdTomato acts a ‘neo-antigen’ when implanted (37). Since the HKP1 tumors grow aggressively in the mice and in most cases do not lose tdTomato, this indicates the tumor normally evades an immune response against it. This is likely to happen, at least partially, through the absence of expression of MHC-I, as indicated in our analyses. This is not unlike what happens in patient tumors which can carry different neo-antigens or tumor associated antigens, such as NY-ESO-1 or MAGE, but downregulate antigen presentation (38). Conversely, we found increased expression of MHC-I and MHC-II on HKP1 cells in F4.CAR-T treated mice, and observed that these remaining cells now presented reduced expression of tdTomato. We and others reported previously this can occur as a consequence of antigen editing, as cancer cells that lose or downregulate antigen are no longer under selective pressure to downregulate the antigen presentation machinery (27,39,40). Our studies therefore suggest that treatment with macrophage-targeted CAR-T promoted T-cell immunity against antigen-expressing HKP1 cell clones and selected for the outgrowth of tumor cells expressing lower amounts of this protein. Though we could not directly measure tdTomato-specific T cells for lack of available MHC tetramers, we did find an expansion of luciferase-specific T cells (28) in the F4.CAR-T treated mice, which further supports the notion that treatment promoted anti-tumor T-cell responses.

We show that release of IFN-γ by F4.CAR-T cells mediated therapeutic benefit in this system and was essential for the immune editing of tdTomato^high^ cancer cells. These results indicate that CAR-T targeting of macrophages resulted in localized IFN-γ release at the tumor site that boosted immune responses against HKP1-associated antigens and immune editing of antigenic tumor cell clones. Interestingly, while Ifng-/- F4.CART cells did not delay tumor growth, they still mediated elimination of target macrophages and eosinophils. This indicates that IFN-γ release was the primary mode of action F4.CAR-T leading to tumor reduction in this setting. This may also be the case for other CAR-T cells that target components of the TME (41). IFN-γ has been established to play key roles in the cancer-immunity cycle and in the treatment efficacy of CAR-T cell therapy (42). Our comparison between wildtype and Ifng-/- F4.CAR-T treated mice strongly suggests that the IFN-γ released from the CAR-T cells was diffusing from the CART-macrophage synapse (43,44) and reprogramming the immunophenotype of the tumor by upregulating MHC-I and MHC-II on tumor and tumor-associated myeloid cells. Though there were likely additional effects of IFN-γ [e.g. turning on the immunoproteosome (45)], upregulated antigen presentation would have promoted priming and increased the sensitivity of cancer cells to cellular immunity. Thus, macrophage-targeted CAR-T may optimally serve as a trojan horse for TME reprogramming to facilitate endogenous tumor immunity.

F4.CAR-T treatment caused an upregulation of PDL1 in tumor and tumor-infiltrating cells in the models tested. HKP1 tumors had been previously shown to be partially responsive to PD1 blockade (29). When tested side-by-side, F4.CAR-T were able to mediate a similar or greater level of tumor reduction as PD1 blockade, but there was no synergistic enhancement of antitumor effects by combining F4.CAR-T cells and PD1 blockade. Notably, we found that PD1 blockade, similar to F4.CAR-T cells, resulted in enrichment of tdTomato^low/neg^ cancer cells. This outgrowth of poorly immunogenic tumor clones could provide an explanation for the ultimate tumor progression in mice treated with either agent, and for the lack of synergy between the two in this model. That is, both immunotherapies induced an anti-tdTomato endogenous immune response which cleared tdTomato^high^ cells, but then tdTomato^low/neg^ cells evaded this response and tumor growth continued. This suggests that a ‘ceiling’ of response may have been reached in the specific models evaluated here which is dependent on the immunogenicity of the cancer cells. Whether this will occur in patients will reflect factors such as the antigenicity of the patient’s tumors.

In addition to lung cancer, we showed that F4.CAR-T treatment can lead to benefit in models of ovarian and pancreatic cancer, and it did so while showing analogous mechanistic traits like the induction of MHC-I expression on tumor cells. Though we did achieve benefit in each model, the F4.CAR-T did not cure animals. As noted above, this appeared to be due to escape by neo-antigen negative/low cells, at least in the HKP1 model. The lack of cure was not unexpected, as our approach did not target cancer cells themselves, and the main rational for developing this approach was to enhance existing treatment strategies. Though in the model we evaluated there was no synergy with PD1 blockade, using a macrophage-targeted CAR-T in conjunction with a cancer cell targeted CAR-T, as Powell and colleagues reported (14), may be the most effective approach. The IFN-γ-dependent induction of MHC-I and II on tumor cells would also suggest that this approach could sensitize tumors to antitumor TCR-based adoptive cell transfer (46), a therapeutic strategy that has already produced some positive results in models analogous to the ones presented here (40,47).

Lymphocyte depletion is typically employed prior to adoptive T cell therapies, including CAR-T cells. This is used to facilitate T cell expansion in the recipient. In our studies, F4.CAR-T cells greatly expanded in mice without the need for prior lymphodepletion. This may have been possible because the cells being targeted, macrophages, are professional antigen presenting cells, which express many immunostimulatory genes, including co-stimulatory molecules and pro-inflammatory cytokines, that can enhance T cell activation. Indeed, two recent studies used mRNA or amph-ligand vaccines to promote expression of a CAR target in dendritic cells and macrophages as a means to expand CAR-T cells in vivo (48,49).

We designed a CAR construct directed against the mouse pan-macrophage marker F4/80 to be able to achieve broad targeting of macrophages. Though clinical translational will be best served with a TAM-specific target, such as FRB, we chose a pan-macrophage approach for proof-of-principle studies in order to probe the safety and efficacy of our approach and gain insight into its mechanisms of action. It is worth noting that broad but transient macrophage depletion may be well tolerated, as demonstrated in studies using pan-targeting drugs such as CSF1/R inhibitors (8). Nonetheless, generation of TAM-specific CARs will undoubtedly provide a better safety profile, and this will require identification of targetable surface proteins that are specific to TAM. FRB is an excellent target, as it is not highly expressed by most tissue-resident macrophages. Though whether it is sufficiently expressed by enough human TAM will need to be established. Recent work from several labs, including our own, have sought to use single cell RNA-seq to identify genes uniquely expressed by TAMs (9,25,50,51). These efforts have resulted in the identification of tumor-enriched macrophage profiles across different cancer types and have identified a number of genes that are relatively restricted to TAMs, such as TREM2 and GPNMB. Though both molecules are constitutively expressed on microglia so their use as a target needs to be considered. Further work could explore whether a threshold level of macrophage density or antigen expression is necessary for therapeutic efficacy (52).

In summary, we have here shown that targeting of macrophages using CAR-T cells can achieve antitumor efficacy as a monotherapy in different models of solid organ tumors, bypassing the need for expression of CAR targets on tumor cells. This approach has promising features to become a new tool in the cancer immunotherapy repertoire, making it possible to deliver CAR-T cells and IFN-γ into solid tumors in a tumor antigen-agnostic manner.

## ACKNOWLEDGMENTS

We thank Ivan Reyes-Torres, Maxime Dhainaut, Abishek Vaidya, Jalal Ahmed and Achuth Nair for helpful discussions. We also thank Alan Soto and Frances Avila, from the Biorepository and Pathology Core at Mount Sinai; and Yu Zhou, from the Biomedical Engineering and Imaging Institute at Mount Sinai, for technical assistance. We thank the Center for Comparative Medicine and Surgery at Mount Sinai for animal care. We also thank Vivek Mittal (Weill Cornell) for the HKP1 cells. B.D.B. was supported by NIH (R01CA257195) and a grant from the Alliance for Cancer Gene Therapy. M.M. was supported by NIH (and R01CA254104) and a grant from the Applebaum Foundation. A.R.S.P. was supported by a grant from Fundación Alfonso Martín Escudero (Spain).

## AUTHOR CONTRIBUTIONS

Conceptualization, A.R.S.P., J.M.T., M.M., B.D.B.; Methodology, A.R.S.P, J.M.T., G.M., S.R.N., A.M., A.L., L.P., A.B.; Writing – Original Draft, A.R.S.P., J.M.T., B.D.B.; Writing – Review & Editing, A.R.S-P., J.M.T., M.M., B.D.B.; Supervision, M.M., B.D.B..; Funding Acquisition: M.M. and B.D.B.

## MATERIALS AND METHODS

### Cloning of CAR construct

F4/80 hybridoma cells were purchased from ATCC (HB-198™). Hybridoma cells were sent to Genscript for Standard Antibody Sequencing for the Variable Domain. The sequence for the identified heavy and light chain portions of the variable domain were synthesized in a ScFv format, separated by a 3x GGGGS linker (GeneArt Gene Synthesis, ThermoFisher). The construct was cloned into an MSGV CD28-CD3z CAR retroviral vector (Addgene #107226). Briefly, an NsiI restriction site was introduced upstream of the backbone’s Kozak region by point mutagenesis, and NsiI and SalI (New England Biolabs) were used to excise the previous CD19 ScFv and introduce the new F4/80 ScFv. PCR overlap assembly (53) was used to introduce a T2A-GFP sequence downstream of CD3z. Co-expression of GFP and the CAR construct was verified by flow cytometry staining of the CAR using an anti-mouse IgG (H+L) antibody (A21235, Invitrogen).

### Cell culture

Phoenix-Eco cells were obtained from ATCC (ATCC CRL-3214™) and cultured in IMDM medium (Gibco 12440-053) supplemented with 10% FBS (Gibco 26140-079), 2 mM Glutamine (Gibco 25030-081) and 1% Pen/Strep (Gibco 15140-122). RAW264.7, A20, HKP1 (24) and mouse CD8 T cells were cultured in RPMI 1640 medium (Gibco 22400-089) containing the previous supplements plus 50 μm b-mercaptoethanol (Gibco 21985-023). ID8_VEGF cells (30) were cultured in RPMI containing the previous supplements plus 500 nM Sodium Pyruvate (Corning 25-000-CI). KPC cells (34) were cultured in DMEM supplemented with 5% FBS (Gibco 26140-079), 2 mM Glutamine (Gibco 25030-081) and 1% Pen/Strep (Gibco 15140-122).

For killing assays, effector : target : non-target cocultures (at an X:1:1 ratio) were cultured for 24h in 96-well culture plates. To minimize alloreactivity-derived target killing, we used Balb/c CD8 CAR-T cells with RAW264.7 macrophages, adding an equal number of A20 B cells as non-target controls. In separate experiments, C57Bl/6 CAR-T cells were cocultured with thioglycate-elicited peritoneal macrophages (see below), using peritoneal neutrophils as non-target controls. 4 hours before the end of the culture, 3 μg/ml Brefeldin A (Invitrogen 00-4506-51) was added to the cultures for subsequent cytokine staining. Specific lysis was calculated as “(*[T/NT]_control_average_-[T/NT]_sample_)/ ([T/NT]_control_average_*100)*”, with T and NT representing the number of target and nontarget cells in each sample.

Generation of thio-elicited macrophages was carried out as previously described (54). Briefly, a 3% solution of Thioglycolate Broth (ThermoFisher CM0173B) was prepared in distilled water, autoclaved and stored at 4°C until use. 1.5 ml of thioglycolate solution was injected intraperitoneally (i.p.) into C57Bl/6 mice. 72h later, mice were euthanized and 5 ml PBS 2mM EDTA injected i.p. to collect the peritoneal lavage. The lavage was RBC-lysed and resuspended in RPMI medium containing 10 ng/ml murine M-CSF (Peprotech 315-02) for co-culture with CAR or control transduced T cells.

### Retroviral vector production

Phoenix-Eco cells were seeded into 10 or 15 cm tissue culture plates 24h prior to achieve an approximate cell density of 70% at the moment of transfection. Transfection was carried out using the calcium-phosphate method. Briefly, CAR or control plasmid constructs were suspended in 0.1x TE and 0.25M CaCl_2_; one volume of 2x HBS was added on a dropwise fashion while continually vortexing and the resulting solution was immediately added onto Phoenix-Eco cells and allowed to sit overnight. IMDM medium was replaced the next morning and supernatants collected and 0.2 μm-filtered 24-30h after that. Supernatant aliquots were stored at −80°C until use. For control purposes throughout all experiments, we used a retroviral HSC1 vector coding for GFP under the EF1α promoter.

### T-cell transduction

CD8 T cells were isolated from mouse spleens using the EasySep™ Mouse CD8+ T Cell Isolation Kit (StemCell 19853). Activation was carried out at a cell density of 1M/ml in 100 U/ml Recombinant Murine IL-2 (Peprotech 212-12). Dynabeads™ Mouse T-Activator CD3/CD28 (ThermoFisher 11453D) were used to activate cells at a 1:4 bead-to-cell ratio for 24h before transduction. Non-treated culture plates (NUNC™, ThermoFisher) were coated overnight at 4°C with 20 μg/ml Retronectin (Takara T100B). Viral supernatant was spun for 90 minutes at 2,000g and 30°C onto the plated retronectin and half the supernatant volume removed carefully after spinning. T cells were resuspended in fresh RPMI medium with IL2, added onto the supernatant-containing wells (final IL-2 concentration 50U/ml), and allowed to sit for 24h. After this, Dynabeads were magnetically removed and T cells were resuspended in fresh RPMI medium at 50 U/ml IL-2. New RPMI medium containing 50 U/ml IL-2 was added daily to keep T cells at a concentration between 1-2.5*10^6^ cells/ml until use, typically within 5 days of isolation. Detection of GFP by flow cytometry was employed to quantify the efficiency of transduction.

### Mouse experimentation

C57Bl/6, CD45.1^+^ C57Bl/6, Ifng-/- C57Bl/6 and Balb/c female mice were obtained from Jackson laboratories (Jackson ID 000664, 002014, 002287 and 000651, respectively) and kept in the Mount Sinai vivarium during use. All mouse experiments were carried out under institutional IACUC approval. All mice were randomized before treatment administration.

For tumor inoculation, HKP1 cells were trypsinized, passed through a 70 μm cell strainer and resuspended in ice-cold sterile PBS at a concentration of 5×10^5^ cells/ml. Mice were heated for 3-5 min using a heating lamp and 1.5×10^5^ cells were injected into the tail vein in a volume of 300 μl using a 27G needle. Anti-PD1 (RMP1-14, BioXCell) was diluted in PBS to a concentration of 1 mg/ml for intraperitoneal administration of 100 ug. Blood samples were drawn where indicated using maxillary vein puncture and stained for flow cytometry. Mice were euthanized at the indicated time points using CO_2_ inhalation. Lungs and spleens were extracted, weighed, and processed for immunofluorescence or flow cytometry.

T cells were injected intravenously via tail vein injection in a volume of 300 μl PBS using 27G needles. The number of T cells to inject in each experimental group was adjusted on the basis of GFP transduction efficiency to have 5×10^6^ GFP^+^ control or F4.CAR-T cells per injection.

For ID8 tumor inoculation, cells were trypsinized and resuspended in PBS at a concentration of 5×10^6^ cells/ml. 10^6^ cells were injected i.p. into female mice. 14 days later, 5×10^6^ CAR-T or control cells were injected i.v. into mice. Mice weight was recorded twice weekly from tumor inoculation. We monitored mice visually for the development of peritoneal ascites and photographed mice at day 24 post-tumor inoculation (representative examples in Fig 6C).

For the experiments involving pancreatic tumors, we used the KPC cell line FC1245 that were established from a tumor in the KPC mouse model as previously described (34). Briefly, female C57Bl/6 mice were anesthetized with ketamine/xylazine. 5×10^4^ tumor cells were suspended in 25 μl matrigel and surgically inoculated into the pancreas. CAR-T cells were administered in 200 μl PBS by retroorbital i.v. injection 7 days later. At day 14 post-tumor inoculation, mice were euthanized and pancreatic tumors obtained, separated from healthy tissue, weighed, and digested in collagenase IV/DNAse I for flow cytometry.

### IVIS

Mice were subjected to anesthesia using inhaled isofluorane and received 100 ul 15 mg/ml luciferase through r.o. injection 5 minutes before image acquisition. Luciferase-based in vivo bioluminescence was acquired in an IVIS Imaging System (Perkin Elmer) at the BioMedical Engineering and Imaging Institute (BMEII) at Mount Sinai. Images were processed using Living Image software (Perkin Elmer).

### Flow cytometry

Lungs were minced and digested for 30 min in Collagenase IV before being mashed through a 70 μm cell strainer. Red blood cells were lysed using RBC Lysis Buffer (eBioscience 00-4300-54). Extracellular stainings were carried out for 10 minutes at 4°C in PBS containing 5% BSA and 2 mM EDTA (Flow Buffer). H2Db-LMYRFEEEL tetramers were generated by the NIH Tetramer Core Facility and allowed to stain at a 1/150 dilution for 30 minutes at 4°C. For intracellular cytokine staining, 3 μg/ml Brefeldin A was added 3-5h before staining. Cells were sequentially stained for extracellular antigens, fixed and permeabilized using BD Cytofix/Cytoperm™ Fixation/Permeabilization Solution Kit before being resuspended in Flow Buffer for acquisition. The cell per organ calculation, where applicable, was carried out using AccuCheck counting beads (Life Technologies). Samples were acquired using FACS Diva in a BD Fortessa flow cytometer and analyzed using FlowJo (BD). The full list of flow antibodies used can be found in Supplementary table 1.

### Immunofluorescence

Lungs were fixed overnight in PLP buffer (in-house) containing 1% PFA, 75 mM L-Lysin and 10 mM NaIO_4_ in pH 7.4 phosphate buffer. After fixation, samples were dehydrated in an increasing sucrose gradient, included in OCT (FisherScientific 23-730-571) and stored at −80°C. 10 μm sections were prepared on a cryostat (Thermo Scientific HM525) and placed on microscopy slides (FisherScientific 12-550-15) for staining. Non-specific binding was blocked using PBS containing 2% FBS and 0.5% BSA. Staining of F4/80 (1/200, Invitrogen MF48021) was carried out overnight at 4°C in PBS containing 1% FBS and 0.25% BSA. DAPI counterstaining (1/5,000, ThermoFisher 62248) was added as a 5 min incubation after antibody binding, and cover slides added for acquisition. Images were obtained at 10x magnification using a Zeiss LSM780 confocal microscope from the Microscopy Core at Mount Sinai. Images were generated using Zeiss Zen software and processed and analyzed using ImageJ and QuPath software. The Cell Detection function within QuPath was used for quantification of macrophages per area of tissue, based on F4/80 staining intensity.

### Histology

The left lobes of mouse lungs were fixed overnight at 4°C in 4% PFA and then immersed in 70% EtOH before processing. Paraffin embedding, generation of 10 μm sections and H&E staining were carried out by the Biorepository and Pathology Core at Mount Sinai. H&E sections were scanned in a Leica Aperio AT2 whole-slide scanner. Quantification of tumor burden on H&E slides was calculated on lung whole slide scans using QuPath software, defining tumor nodules by morphological features. Histological tumor burden was calculated as the sum of the areas of all tumor nodules divided by the total lung area and expressed as a percentage.

### Statistical analyses

Statistical analyses were carried out using GraphPad Prism 9 software. Log-rank analysis was used to compare survival curves. T-student comparison was used for column analyses, except where indicated: Mann-Whitney rank-based analyses were used to include placeholder data points representing mice that succumbed to tumor development before sacrifice (indicated in figures with an “x” datapoint symbol).

### Graphical resources

BioRender was used for the elaboration of the graphical abstract and illustrative guides.

---

**Supplementary Figure 1.**
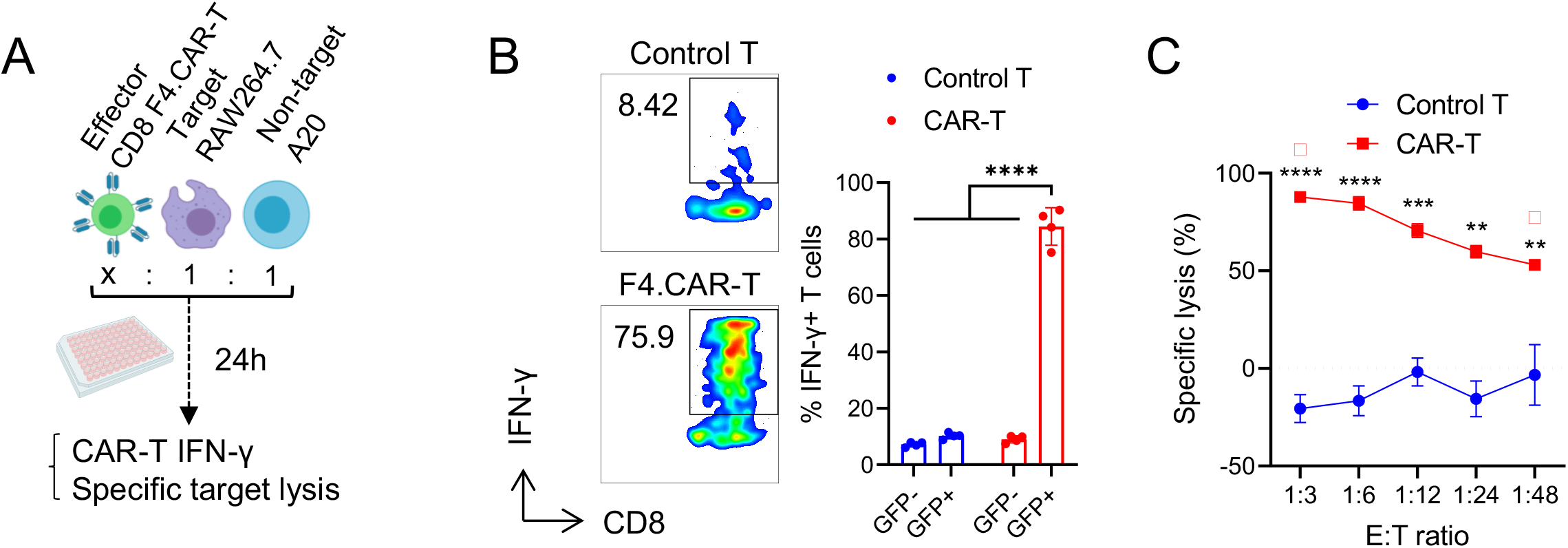
F4/80 CAR-T cells eliminate RAW264.7 macrophages in vitro. **(A)** Schematic explaining the assay design. Numbers below the indicated cells indicate starting relative cell numbers. **(B)** On the left, representative flow cytometry plots showing intracellular IFN-γ staining on GFP+ control or CAR-T cells. Shown are the percentages of IFN-γ^+^ cells. On the right, quantification of IFN-γ^+^ cells across replicates. **(C)** Specific lysis of RAW264.7 cells across a range of E:T ratios. *p<0.05, **p<0.01, ***p<0.001, ****p p<0.0001

**Supplementary Figure 2.**
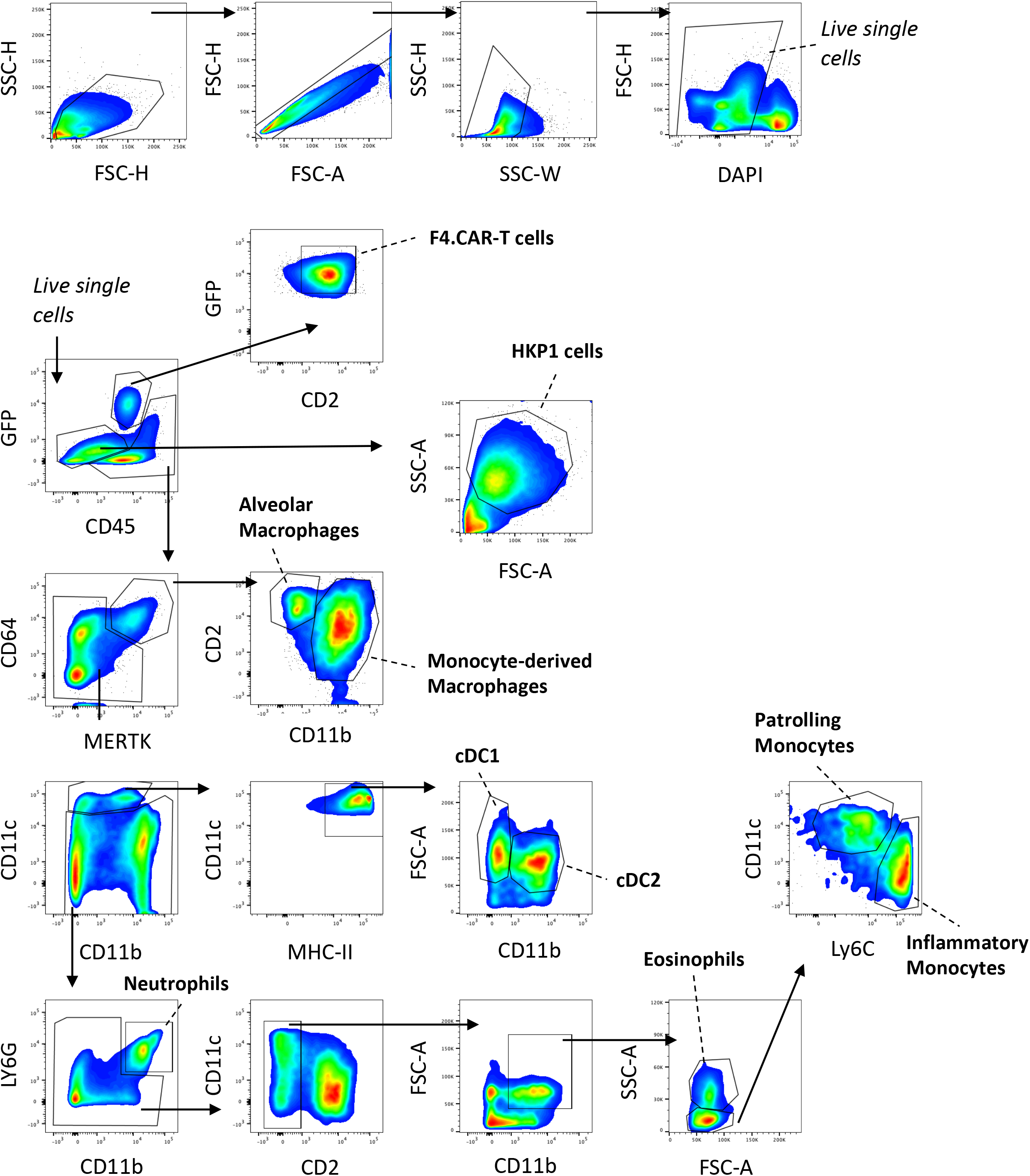
Flow cytometry gating strategy in HKP1-bearing lungs. Depicted is a representative flow cytometry gating for an untreated lung bearing HKP1 tumors. Plots from an F4.CART-treated mouse (GFP vs CD45; GFP vs CD2) are added to demonstrate CAR-T gating.

**Supplementary Figure 3.**
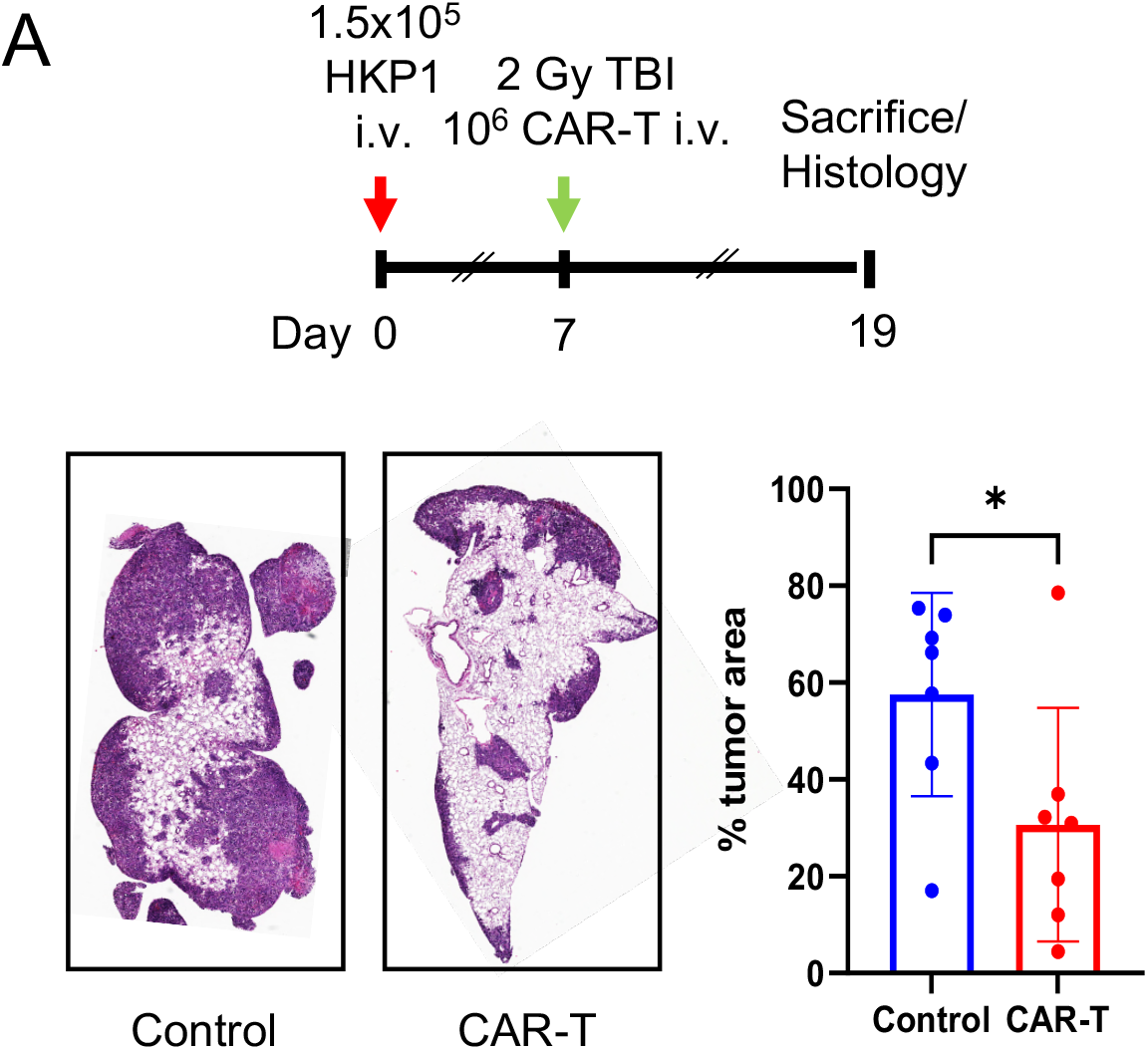
F4.CAR-T delays growth of HKP1 tumors following lymphodepleting preconditioning. HKP1 tumors were established and CAR-T cells administered according to the treatment schematic. Mice were euthanized at the indicated time point and lungs processed for H&E staining. Left panels show representative sections from control or F4.CAR-T treated mice. Histological tumor burden was calculated as described in materials and methods. *p<0.05

**Supplementary Figure 4.**
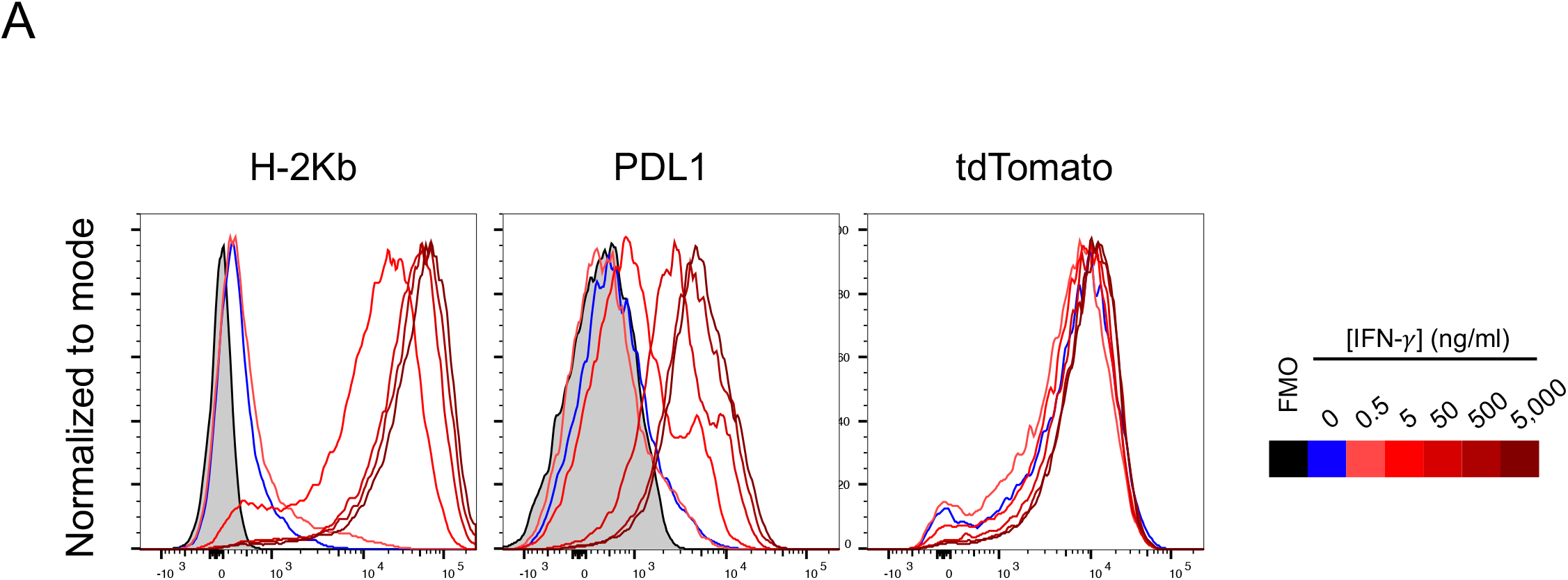
IFNg does not directly affect tdTomato expression by HKP1 cells. HKP1 cells were exposed in vitro to recombinant murine IFNg (Peprotech) at the listed concentrations for 24h. After this, cells were trypsinized and stained for MHC-I and PDL1. Shown are representative histograms for MHC-I, PDL1 and tdTomato detection on live HKP1 cells. Histogram color legend on the right. FMO: field-minus-one for MHC-I or PDL1 stainings.

**Supplementary Figure 5.**
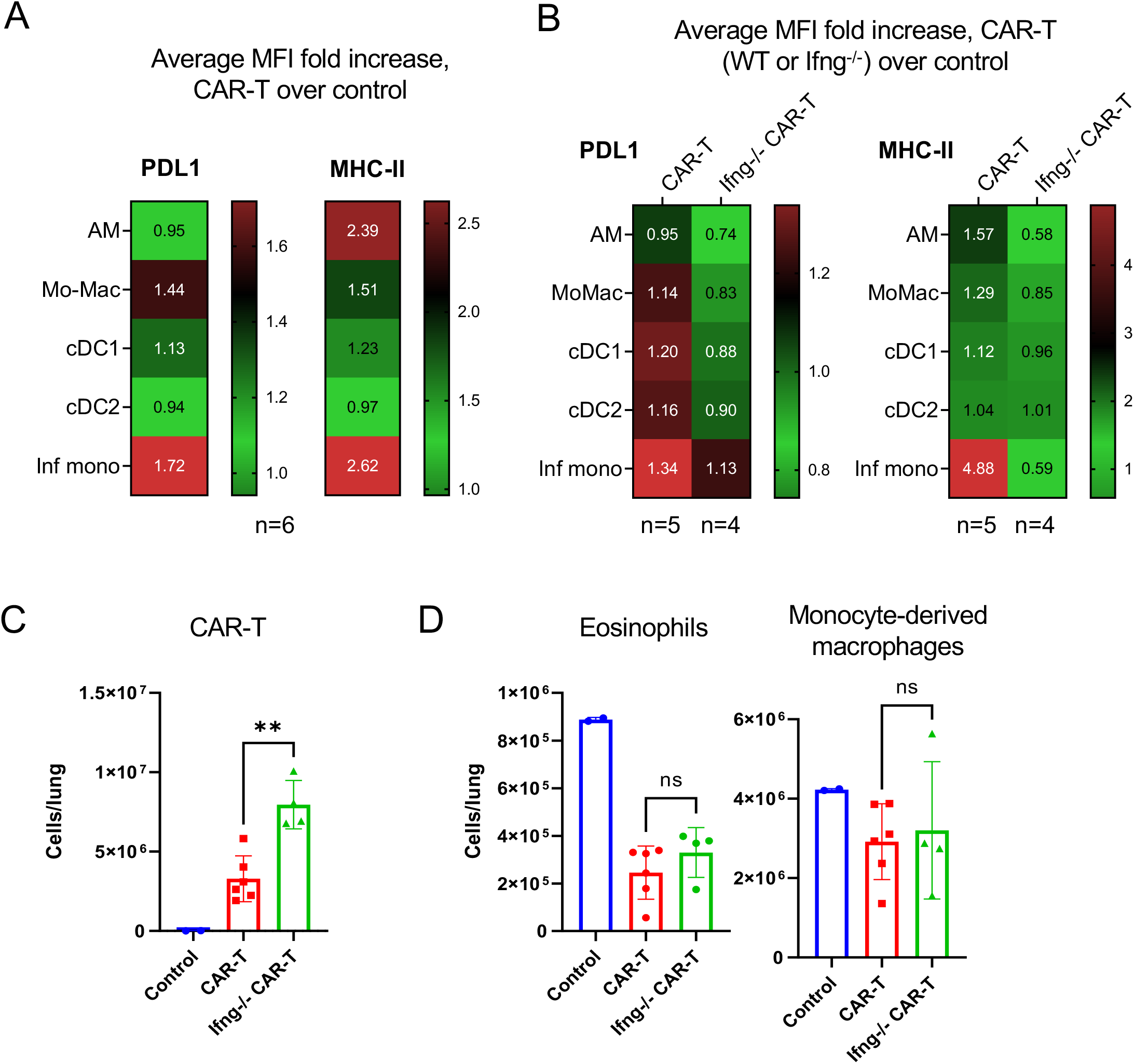
IFN-*γ* release by F4.CAR-T cells reprograms the tumor myeloid microenvironment but is not required for target cell lysis. HKP1 tumors were established and treated as in Supp.Fig.3B (A) or Fig 5 (B-D). Lung samples were processed for FC and stained. **(A,B)** MFI were calculated for PDL1 and MHC-II for the listed myeloid populations. Shown are the average fold changes in these populations in F4.CAR-T treated mice over controls. **(C,D)** Shown are the number of F4.CAR-T cells (C) and eosinophils and MoMacs **(D)** in the lungs of control, F4.CAR-T or Ifng-/- F4.CAR-T treated mice. AM: alveolar macrophage; Mo-Mac: monocyte-derived macrophage; cDC1/2: conventional dendritic cell 1/2; Inf mono: inflammatory monocyte. *p<0.05, **p<0.01, ***p<0.001, ****p p<0.0001

**Supplementary table 1.**
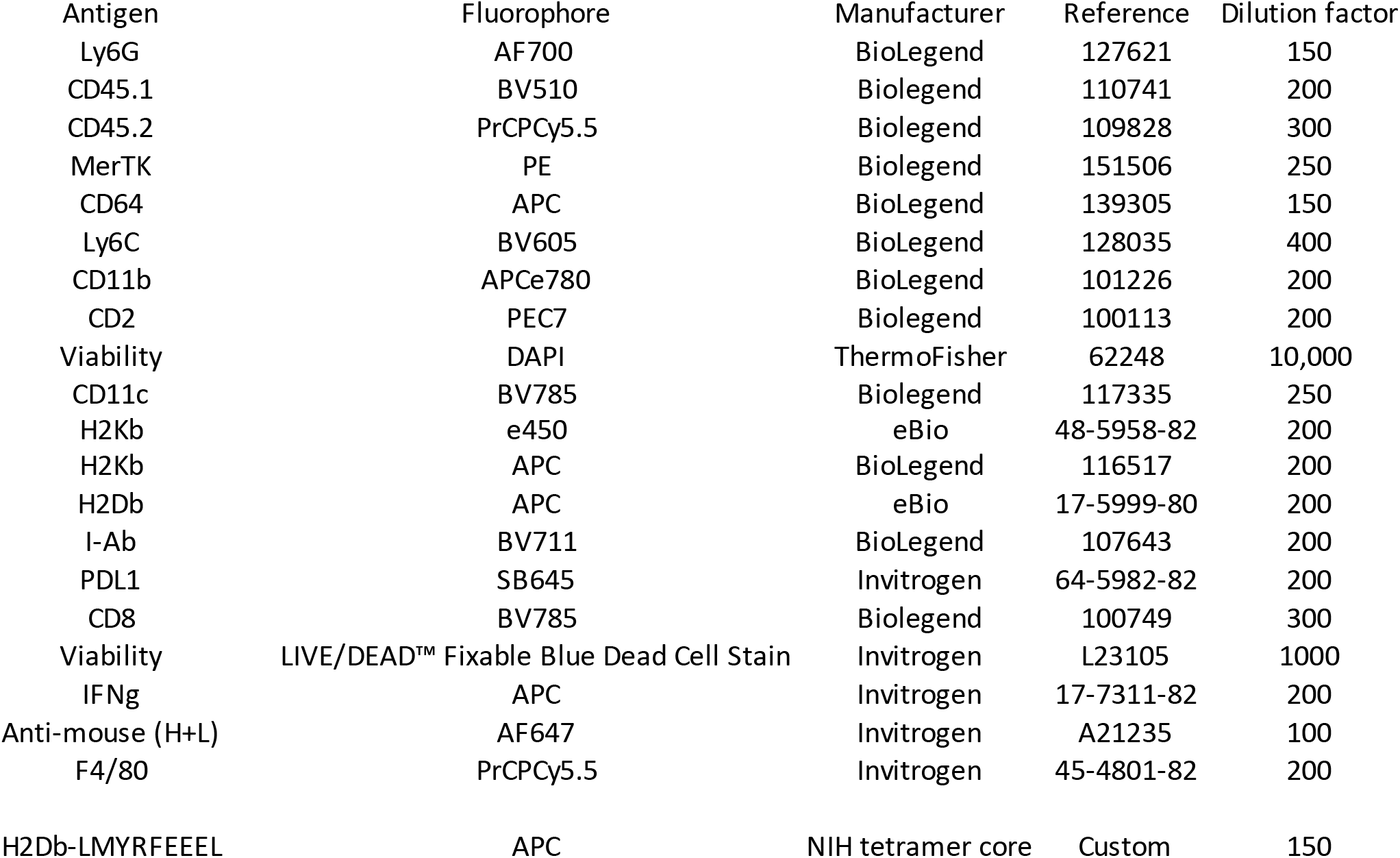
Flow cytometry reagents

## REFERENCES

1. Hegde PS, Chen DS. Top 10 Challenges in Cancer Immunotherapy. Immunity. 2020;52:17–35.

2. Mantovani A, Marchesi F, Malesci A, Laghi L, Allavena P. Tumour-associated macrophages as treatment targets in oncology. Nat Rev Clin Oncol. Nature Publishing Group; 2017;14:399–416.

3. Binnewies M, Roberts EW, Kersten K, Chan V, Fearon DF, Merad M, et al. Understanding the tumor immune microenvironment (TIME) for effective therapy. Nat Med. Springer US; 2018;24:541–50.

4. Noy R, Pollard JW. Tumor-Associated Macrophages: From Mechanisms to Therapy. Immunity. Elsevier Inc.; 2014;41:49–61.

5. DeNardo DG, Ruffell B. Macrophages as regulators of tumour immunity and immunotherapy. Nat Rev Immunol. Springer US; 2019;19:369–82.

6. Cassetta L, Pollard JW. Targeting macrophages: Therapeutic approaches in cancer. Nat Rev Drug Discov. Nature Publishing Group; 2018;17:887–904.

7. Gomez-Roca CA, Cassier PA, Italiano A, Cannarile M, Ries C, Brillouet A, et al. Phase I study of RG7155, a novel anti-CSF1R antibody, in patients with advanced/metastatic solid tumors. J Clin Oncol. 2015;33:3005.

8. Cannarile MA, Weisser M, Jacob W, Jegg A-MM, Ries CH, Rüttinger D. Colony-stimulating factor 1 receptor (CSF1R) inhibitors in cancer therapy. J Immunother Cancer. Journal for ImmunoTherapy of Cancer; 2017;5:1–13.

9. Molgora M, Esaulova E, Vermi W, Hou J, Chen Y, Luo J, et al. TREM2 Modulation Remodels the Tumor Myeloid Landscape Enhancing Anti-PD-1 Immunotherapy. Cell. Elsevier; 2020;182:886–900.e17.

10. Binnewies M, Pollack JL, Rudolph J, Dash S, Abushawish M, Lee T, et al. Targeting TREM2 on tumor-associated macrophages enhances immunotherapy. Cell Rep. 2021;37:109844.

11. Zhang F, Parayath NN, Ene CI, Stephan SB, Koehne AL, Coon ME, et al. Genetic programming of macrophages to perform anti-tumor functions using targeted mRNA nanocarriers. Nat Commun 2019 101. Nature Publishing Group; 2019;10:1–16.

12. Ruella M, Klichinsky M, Kenderian SS, Shestova O, Ziober A, Kraft DO, et al. Overcoming the Immunosuppressive Tumor Microenvironment of Hodgkin Lymphoma Using Chimeric Antigen Receptor T Cells. Cancer Discov. 2017;7:1154–67.

13. Parihar R, Rivas CH, Huynh M, Omer B, Lapteva N, Metelitsa LS, et al. NK cells expressing a chimeric activating receptor eliminate MDSCs and rescue impaired CAR-T cell activity against solid tumors. Cancer Immunol Res. 2019;7:363–75.

14. Rodriguez-Garcia A, Lynn RC, Poussin M, Eiva MA, Shaw LC, O’Connor RS, et al. CART cell-mediated depletion of immunosuppressive tumor-associated macrophages promotes endogenous antitumor immunity and augments adoptive immunotherapy. Nat Commun. Nature Publishing Group; 2021;12:877.

15. June CH, Sadelain M. Chimeric Antigen Receptor Therapy. N Engl J Med. 2018;379:64–73.

16. Maude SL, Frey N, Shaw PA, Aplenc R, Barrett DM, Bunin NJ, et al. Chimeric Antigen Receptor T Cells for Sustained Remissions in Leukemia. N Engl J Med. 2014;371:1507–17.

17. Mikkilineni L, Kochenderfer JN. CAR T cell therapies for patients with multiple myeloma. Nat Rev Clin Oncol. 2021;18:71–84.

18. Eyquem J, Mansilla-Soto J, Giavridis T, Van Der Stegen SJC, Hamieh M, Cunanan KM, et al. Targeting a CAR to the TRAC locus with CRISPR/Cas9 enhances tumour rejection. Nature. Nature Publishing Group; 2017;543:113–7.

19. Hong M, Clubb JD, Chen YY. Engineering CAR-T Cells for Next-Generation Cancer Therapy. Cancer Cell. Elsevier Inc.; 2020;38:473–88.

20. Rodriguez-Garcia A, Palazon A, Noguera-Ortega E, Powell DJ, Guedan S. CAR-T Cells Hit the Tumor Microenvironment: Strategies to Overcome Tumor Escape. Front Immunol. 2020;11:1–17.

21. Gordon S, Hamann J, Lin H, Stacey M. F4/80 and the related adhesion-GPCRs. Eur J Immunol. 2011;41:2472–6.

22. Austyn JM, Gordon S. F4/80, a monoclonal antibody directed specifically against the mouse macrophage. Eur J Immunol. 1981;11:805–15.

23. Kochenderfer JN, Yu Z, Frasheri D, Restifo NP, Rosenberg SA. Adoptive transfer of syngeneic T cells transduced with a chimeric antigen receptor that recognizes murine CD19 can eradicate lymphoma and normal B cells. Blood. 2010;116:3875–86.

24. Choi H, Sheng J, Gao D, Li F, Durrans A, Ryu S, et al. Transcriptome Analysis of Individual Stromal Cell Populations Identifies Stroma-Tumor Crosstalk in Mouse Lung Cancer Model. Cell Rep. The Authors; 2015;10:1187–201.

25. Casanova-Acebes M, Dalla E, Leader AM, LeBerichel J, Nikolic J, Morales BM, et al. Tissue-resident macrophages provide a pro-tumorigenic niche to early NSCLC cells. Nature. 2021;595:578–84.

26. Leader AM, Grout JA, Maier BB, Nabet BY, Park MD, Tabachnikova A, et al. Single-cell analysis of human non-small cell lung cancer lesions refines tumor classification and patient stratification. Cancer Cell. Cell Press; 2021;

27. Wroblewska A, Dhainaut M, Ben-Zvi B, Rose SA, Park ES, Amir E-AAD, et al. Protein Barcodes Enable High-Dimensional Single-Cell CRISPR Screens. Cell. Elsevier Inc.; 2018;175:1141–1155.e16.

28. Limberis MP, Bell CL, Wilson JM. Identification of the murine firefly luciferase-specific CD8 T-cell epitopes. Gene Ther. 2009;16:441–7.

29. Markowitz GJ, Havel LS, Crowley MJP, Ban Y, Lee SB, Thalappillil JS, et al. Immune reprogramming via PD-1 inhibition enhances early-stage lung cancer survival. JCI insight. 2018;3:e96836.

30. Hartl CA, Bertschi A, Puerto RB, Andresen C, Cheney EM, Mittendorf EA, et al. Combination therapy targeting both innate and adaptive immunity improves survival in a pre-clinical model of ovarian cancer. J Immunother Cancer. 2019;7:199.

31. Etzerodt A, Moulin M, Doktor TK, Delfini M, Mossadegh-Keller N, Bajenoff M, et al. Tissue-resident macrophages in omentum promote metastatic spread of ovarian cancer. J Exp Med. 2020;217:e20191869.

32. Vonderheide RH, Bear AS. Tumor-Derived Myeloid Cell Chemoattractants and T Cell Exclusion in Pancreatic Cancer. Front Immunol. 2020;11:1–8.

33. Stromnes IM, Hulbert A, Pierce RH, Greenberg PD, Hingorani SR. T-cell Localization, Activation, and Clonal Expansion in Human Pancreatic Ductal Adenocarcinoma. Cancer Immunol Res. Cancer Immunol Res; 2017;5:978–91.

34. Boj SF, Hwang C Il, Baker LA, Chio IIC, Engle DD, Corbo V, et al. Organoid models of human and mouse ductal pancreatic cancer. Cell. sciencedirect; 2015;160:324–38.

35. Connor AA, Denroche RE, Jang GH, Lemire M, Zhang A, Chan-Seng-Yue M, et al. Integration of Genomic and Transcriptional Features in Pancreatic Cancer Reveals Increased Cell Cycle Progression in Metastases. Cancer Cell. 2019;35:267–282.e7.

36. DuPage M, Dooley AL, Jacks T. Conditional mouse lung cancer models using adenoviral or lentiviral delivery of Cre recombinase. Nat Protoc. Nat Protoc; 2009;4:1064–72.

37. Huang L, Bommireddy R, Munoz LE, Guin RN, Wei C, Ruggieri A, et al. Expression of tdTomato and luciferase in a murine lung cancer alters the growth and immune microenvironment of the tumor. PLoS One. 2021;16:e0254125.

38. Dhatchinamoorthy K, Colbert JD, Rock KL. Cancer Immune Evasion Through Loss of MHC Class I Antigen Presentation. Front Immunol. Frontiers Media S.A.; 2021;12:469.

39. Fitzgerald B, Connolly KA, Cui C, Fagerberg E, Mariuzza DL, Hornick NI, et al. A mouse model for the study of anti-tumor T cell responses in Kras-driven lung adenocarcinoma. Cell Reports Methods. Cell Press; 2021;1:100080.

40. DuPage M, Cheung AF, Mazumdar C, Winslow MM, Bronson R, Schmidt LM, et al. Endogenous T cell responses to antigens expressed in lung adenocarcinomas delay malignant tumor progression. Cancer Cell. Elsevier Inc.; 2011;19:72–85.

41. Bughda R, Dimou P, D’Souza RR, Klampatsa A. Fibroblast Activation Protein (FAP) - Targeted CAR-T Cells: Launching an Attack on Tumor Stroma. ImmunoTargets Ther. Dove Press; 2021;Volume 10:313–23.

42. Alizadeh D, Wong RA, Gholamin S, Maker M, Aftabizadeh M, Yang X, et al. IFNγ is Critical for CAR T Cell–mediated Myeloid Activation and Induction of Endogenous Immunity. Cancer Discov. 2021;11:2248–65.

43. Hoekstra ME, Bornes L, Dijkgraaf FE, Philips D, Pardieck IN, Toebes M, et al. Longdistance modulation of bystander tumor cells by CD8+ T-cell-secreted IFN-γ. Nat Cancer. Springer Science and Business Media LLC; 2020;1:291–301.

44. Thibaut R, Bost P, Milo I, Cazaux M, Lemaître F, Garcia Z, et al. Bystander IFN-γ activity promotes widespread and sustained cytokine signaling altering the tumor microenvironment. Nat Cancer. Springer Science and Business Media LLC; 2020;1:302–14.

45. Aki M, Shimbara N, Takashina M, Akiyama K, Kagawa S, Tamura T, et al. Interferon-γ Induces Different Subunit Organizations and Functional Diversity of Proteasomes. J Biochem. Oxford Academic; 1994;115:257–69.

46. Hinrichs CS, Rosenberg SA. Exploiting the curative potential of adoptive T-cell therapy for cancer. Immunol Rev. John Wiley & Sons, Ltd; 2014;257:56–71.

47. Anderson KG, Voillet V, Bates BM, Chiu EY, Burnett MG, Garcia NM, et al. Engineered Adoptive T-cell Therapy Prolongs Survival in a Preclinical Model of Advanced-Stage Ovarian Cancer. Cancer Immunol Res. American Association for Cancer Research; 2019;7:1412–25.

48. Reinhard K, Rengstl B, Oehm P, Michel K, Billmeier A, Hayduk N, et al. An RNA vaccine drives expansion and efficacy of claudin-CAR-T cells against solid tumors. Science (80-). 2020;367:446–53.

49. Ma L, Dichwalkar T, Chang JYHH, Cossette B, Garafola D, Zhang AQ, et al. Enhanced CAR–T cell activity against solid tumors by vaccine boosting through the chimeric receptor. Science (80-). 2019;365:162–8.

50. Zilionis R, Engblom C, Pfirschke C, Savova V, Zemmour D, Saatcioglu HD, et al. Single-Cell Transcriptomics of Human and Mouse Lung Cancers Reveals Conserved Myeloid Populations across Individuals and Species. Immunity. Elsevier Inc.; 2019;50:1317–1334.e10.

51. Mulder K, Patel AA, Kong WT, Piot C, Halitzki E, Dunsmore G, et al. Cross-tissue singlecell landscape of human monocytes and macrophages in health and disease. Immunity. Elsevier; 2021;54:1883–1900.e5.

52. Majzner RG, Rietberg SP, Sotillo E, Dong R, Vachharajani VT, Labanieh L, et al. Tuning the Antigen Density Requirement for CAR T-cell Activity. Cancer Discov. American Association for Cancer Research (AACR); 2020;10:702–23.

53. Bryksin A V., Matsumura I. Overlap extension PCR cloning: A simple and reliable way to create recombinant plasmids. Biotechniques. Eaton Publishing Company; 2010;48:463–5.

54. Turnbull IR, Gilfillan S, Cella M, Aoshi T, Miller M, Piccio L, et al. Cutting Edge: TREM-2 Attenuates Macrophage Activation. J Immunol. 2006;177:3520–4.

